# Differential effects of HDAC inhibitors in the *Rho*^I255d^ mouse model for autosomal dominant retinitis pigmentosa

**DOI:** 10.1101/2025.07.25.666875

**Authors:** Yu Zhu, Pranav Nanda Kumar, Kangwei Jiao, François Paquet-Durand

## Abstract

Retinitis Pigmentosa (RP) is an inherited neurodegenerative disease which leads to loss of retinal photoreceptors and blindness. Histone deacetylases (HDAC) were previously found to be involved in photoreceptor cell death, and HDAC inhibitors have shown protective effects in animal models for autosomal recessive RP. However, whether HDAC inhibitors can protect photoreceptors in autosomal dominant RP (ADRP) remains unclear. Here, we utilized the recently generated human homologous *Rho*^I255d/+^ ADRP mouse model to investigate degenerative mechanisms and the therapeutic potential of HDAC inhibitors.

To visualize photoreceptor HDAC activity, we applied an *in situ* HDAC activity assay on post-natal (P) day 20 wild type (WT) and *Rho*^I255d/+^ retina. Treatment with the HDAC class I/II inhibitor Trichostatin A and the HDAC class III inhibitor nicotinamide (NAM) suggested that most HDAC activity detected in *Rho*^I255d/+^ photoreceptors was related to class I/II isoforms.

The therapeutic potential of different HDAC inhibitors, targeting different HDAC isoforms, was evaluated on organotypic retinal explants cultured under completely controlled conditions. HDAC inhibitors tested included SAHA (Vorinostat), MPT0G211, ACY-957, and NAM. Readouts comprised the TUNEL assay, immunostaining for activated calpain-2 and caspase-3, cone arrestin-3, and bromodeoxyuridine (BrdU)-labeling. Among the compounds tested, MPT0G211, targeting predominantly cytoplasmic HDAC-6, exhibited the strongest protective effect on both rod and cone photoreceptors. Remarkably, ACY-957, inhibiting nuclear HDAC-1/-2, at higher concentration induced both photoreceptor cell death and cell proliferation. High levels of NAM, blocking mitochondrial and nuclear HDACs, caused selective rod cell death, without affecting cones. All HDAC inhibitors tested had no or only minor effects on neurons of the inner retina.

Our study highlights the complexity of HDAC activity during photoreceptor neurodegeneration and cautions against the use of unspecific inhibitors. At the same time, it showcases important differences between rod and cone photoreceptors and suggests especially HDAC-6 as a potential target for future therapy development.

**Graphical abstract:** 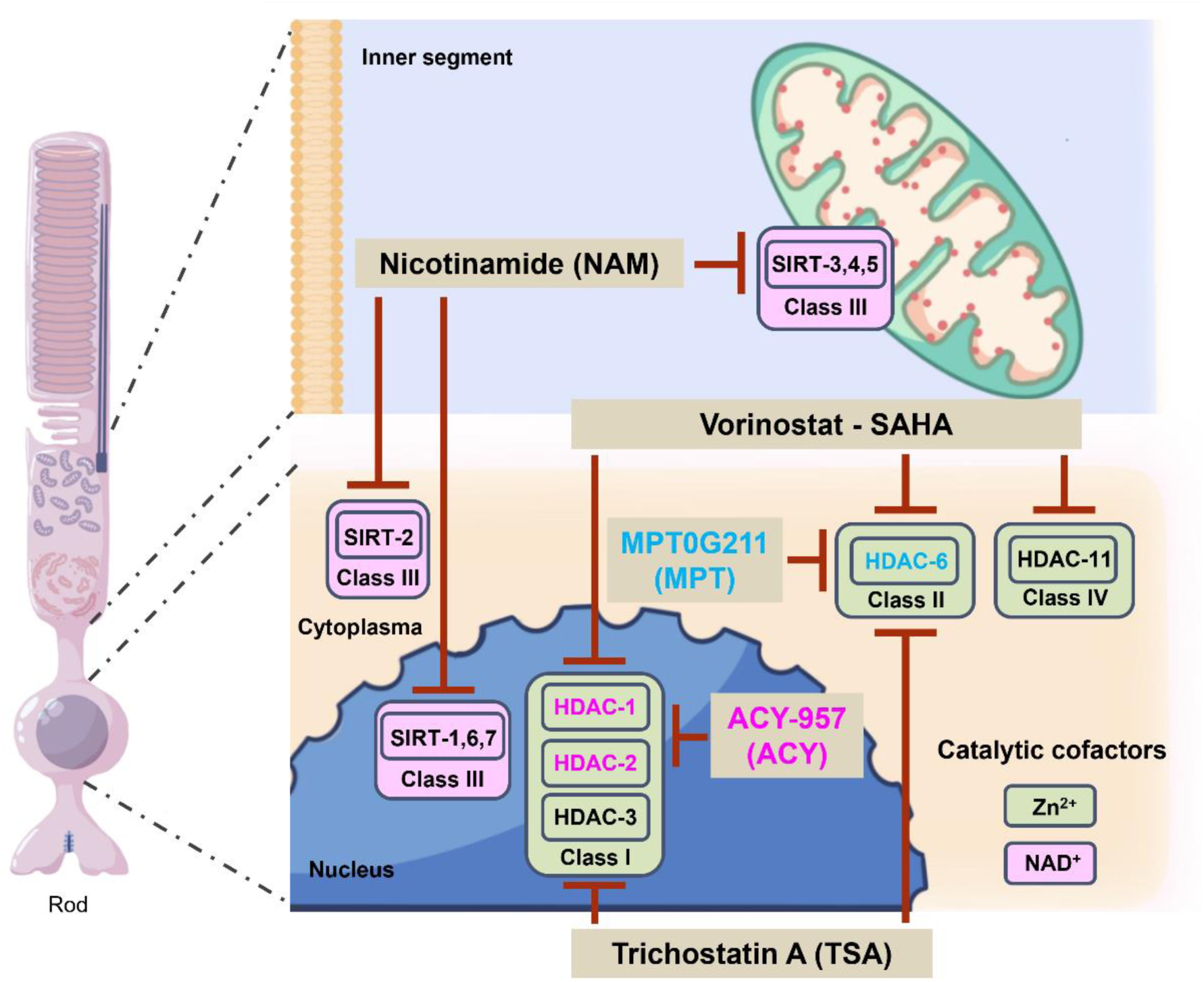

## Introduction

Retinitis pigmentosa (RP) relates to a diverse group of inherited retinal diseases (IRD) characterized by progressive loss of rod photoreceptors, secondary cone photoreceptor cell death, and eventually complete blindness (1). RP remains essentially untreatable, and therapy development is hampered by strong genetic heterogeneity and varied inheritance patterns. The latter may broadly be categorized into autosomal dominant (ADRP), autosomal recessive (ARRP), and X-linked (2–4). About 25% of ADRP cases are caused by rhodopsin (*RHO*) gene mutations (1). Previous studies, using IRD models such as *rd1* and *rd10* mice for ARRP, or *Rho*^S334ter^ and *Rho*^P23H^ ADRP rats (5), found photoreceptor death to associate with heightened activity of histone deacetylases (HDACs). HDACs mediate protein deacetylation in mitochondria, cytoplasm, and nucleus, and regulate gene transcription (5, 6). However, excessive HDAC activity has also been connected to neuronal cell death (7), including in the retina (5, 6). The 18 known HDAC isoforms are grouped into four classes (I-IV) of which class I, II, and IV are zinc-dependent metalloenzymes, while class III is nicotinamide adenine dinucleotide (NAD^+^)-dependent (8) and referred to as sirtuin-type HDACs. Inhibition of HDACs improved photoreceptor survival in various IRD models. Trichostatin A (TSA), an inhibitor for HDAC class I, II, and IV (9, 10), promoted cone photoreceptor survival (11–15). Similarly, the pan-HDAC inhibitor suberoylanilide hydroxamic acid (SAHA; aka Vorinostat), protected rod photoreceptors in *rd1* and *rd10* mice (14, 16).

The recently generated human homologous *Rho*^I255d/+^ knock-in mouse model for ADRP displays a rapid loss of rod and cone photoreceptors, associated with both apoptotic and non-apoptotic, cGMP-dependent degenerative pathways (17, 18), similar to what was previously observed in the transgenic *Rho*^S334ter^ rat model for ADRP (5).

Earlier studies found promising neuroprotective effects of HDAC inhibitors in ARRP models, yet their efficacy in ADRP is still largely unknown. Here, we used the new human homologous *Rho*^I255d^ mouse model for ADRP (17, 18) to investigate the role of HDACs in photoreceptor cell death and the potential therapeutic effects of HDAC inhibitors. We found that, while HDAC inhibition can delay progression of photoreceptor loss, specific HDAC inhibitors can also have strong detrimental effects. This data expands our understanding of ADRP pathogenesis and may advance development of new treatment options for ADRP.

## Results

### HDAC class I/II activity increases in *Rho*^I255d/+^ retina

To assess HDAC activity in *Rho*^I255d/+^ retina, we employed an *in situ* HDAC activity assay (15) on fixed retinal sections derived from wild type (WT) and *Rho*^I255d/+^ mice at post-natal (P) day 20. To distinguish between the major HDAC classes, we first used the well-established pan-inhibitors trichostatin A (TSA; class I/II) and nicotinamide (NAM; class III), or both in combination (19–23) (for compound specificity see Table 1).

**Table 1:**
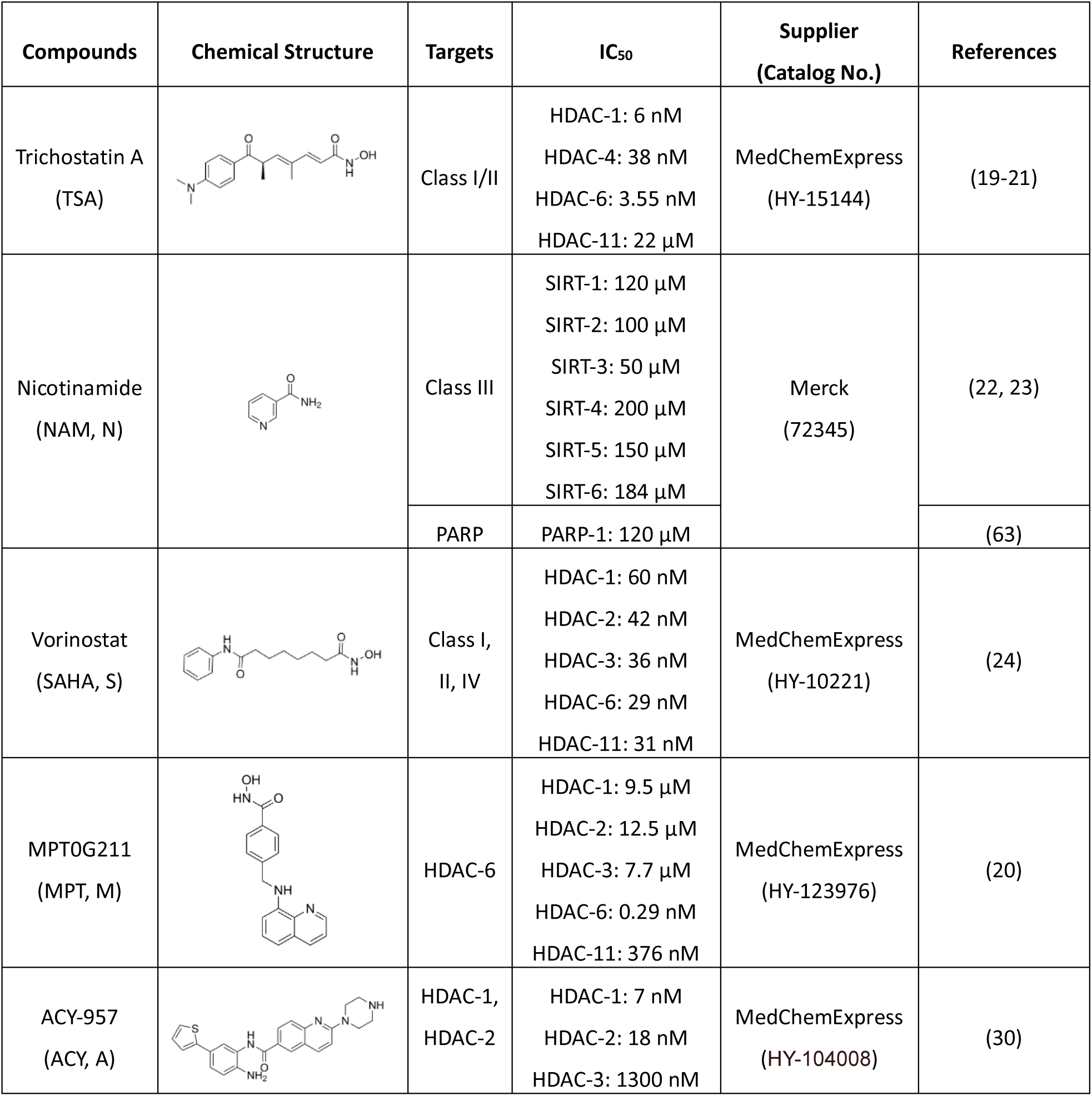
Overview of different HDAC inhibitors used. Chemical structures, targets, IC_50_, suppliers, and references.

| Compounds | Chemical Structure | Targets | IC<sub>50</sub> | Supplier<br>(Catalog No.) | References |
| --- | --- | --- | --- | --- | --- |
| Trichostatin A<br>(TSA) |  | Class I/II | HDAC-1: 6 nM<br>HDAC-4: 38 nM<br>HDAC-6: 3.55 nM<br>HDAC-11: 22 μM | MedChemExpress<br>(HY-15144) | (19-21) |
| Nicotinamide<br>(NAM, N) |  | Class III | SIRT-1: 120 μM<br>SIRT-2: 100 μM<br>SIRT-3: 50 μM<br>SIRT-4: 200 μM<br>SIRT-5: 150 μM<br>SIRT-6: 184 μM | Merck<br>(72345) | (22, 23) |
|  |  | PARP | PARP-1: 120 μM |  | (63) |
| Vorinostat<br>(SAHA, S) |  | Class I,<br>II, IV | HDAC-1: 60 nM<br>HDAC-2: 42 nM<br>HDAC-3: 36 nM<br>HDAC-6: 29 nM<br>HDAC-11: 31 nM | MedChemExpress<br>(HY-10221) | (24) |
| MPTOG211<br>(MPT, M) |  | HDAC-6 | HDAC-1: 9.5 μM<br>HDAC-2: 12.5 μM<br>HDAC-3: 7.7 μM<br>HDAC-6: 0.29 nM<br>HDAC-11: 376 nM | MedChemExpress<br>(HY-123976) | (20) |
| ACY-957<br>(ACY, A) |  | HDAC-1,<br>HDAC-2 | HDAC-1: 7 nM<br>HDAC-2: 18 nM<br>HDAC-3: 1300 nM | MedChemExpress<br>(HY-104008) | (30) |

HDAC activity was increased in *Rho*^I255d/+^ outer nuclear layer (ONL; *i.e.* the photoreceptor layer) compared to WT (Fig. 1A, C). In WT retina, treatment with TSA, NAM, or both compounds combined (Fig. 1A, B) did not significantly change the numbers of HDAC-positive cells (Fig. 1E). However, in the *Rho*^I255d/+^ mutant situation, non-treated (NT) retinas showed large numbers of HDAC activity positive cells compared to retinas that had been treated with either TSA, NAM, or a combination of both (Fig. 1C, D). TSA treatment reduced HDAC activity in the ONL even more strongly than NAM (Fig. 1E), indicating that in *Rho*^I255d/+^ photoreceptors, excessive activation of HDAC was primarily dependent on HDAC isoforms belonging to class I/II (Table S1).

**Figure 1.**
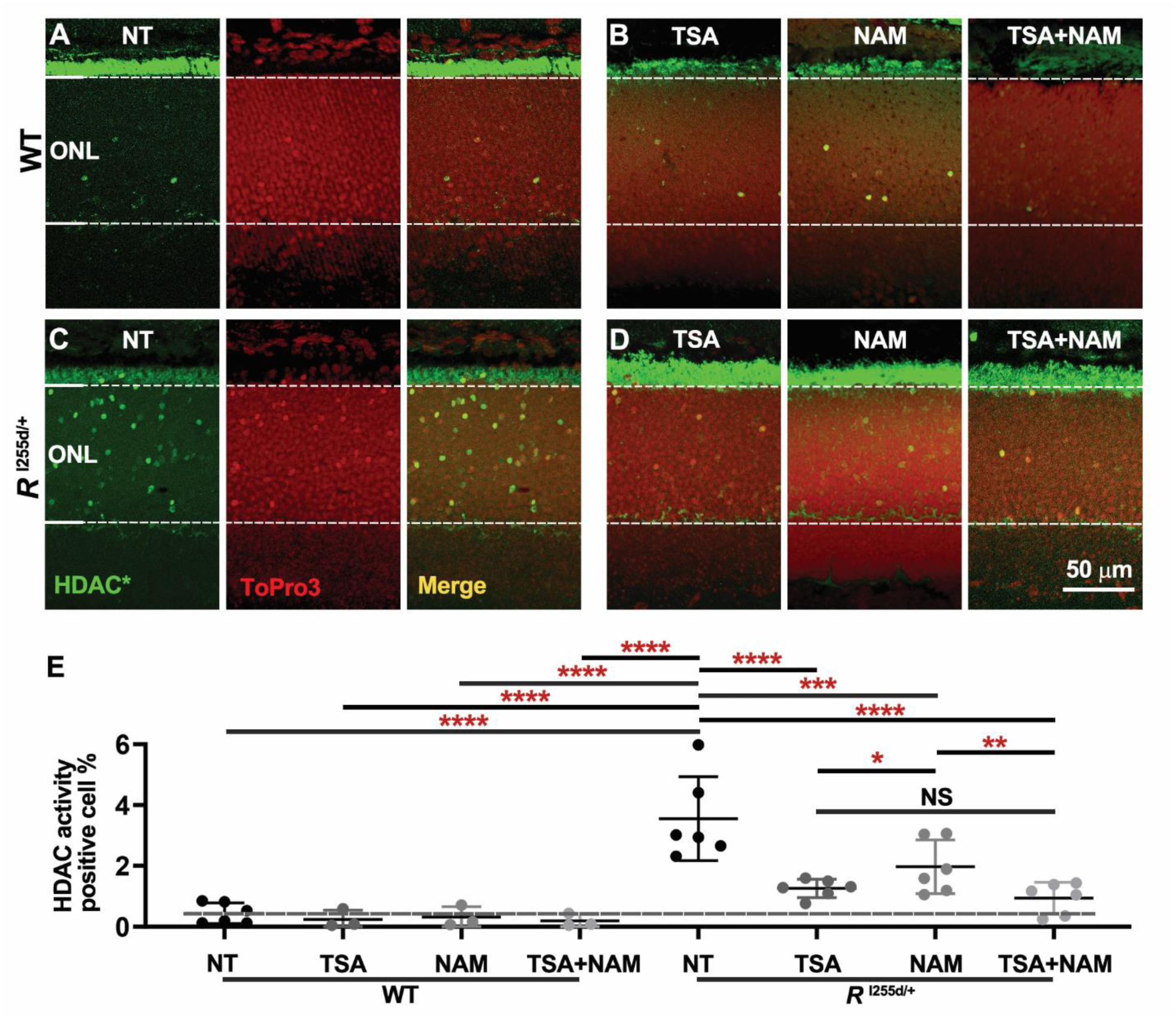
Increased HDAC activity in *Rho*^I255d/+^ mutant retina. A-D. Retinal cryosections derived from wild-type (WT) and *Rho*^I255d/+^ mice at post-natal day 20 were stained for HDAC activity (HDAC*, green). **A, C.** Compared to WT, the outer nuclear layer (ONL) of *Rho*^I255d/+^ mutant retina displayed increased numbers of cells with HDAC*. ToPro3 (red) was used as nuclear counterstain, the 3^rd^ image in a segment shows merged green and red channels. **B, D.** HDACs belonging to class I/II were inhibited with TSA, while class III was blocked with NAM. TSA and NAM combined were used to block most HDAC*. **E.** Quantification of HDAC*-positive cells in ONL. HDAC inhibitor treatment did not significantly change the number of HDAC*-positive cells in WT retina. High numbers of HDAC*-positive cells were observed in non-treated (NT) mutant ONL. Both TSA and NAM reduced HDAC* in *Rho*^I255d/+^ retina, with the combined treatment displaying the lowest positive cell counts. Images are representative of results obtained from 3-6 independent specimens; data points represent mean and SD; statistical analysis: Two-way ANOVA with two-stage linear step-up procedure of Benjamini, Krieger, and Yekutieli; * = *p* ≤ 0.05, ** = *p* ≤ 0.01, *** = *p* ≤ 0.001, **** = *p* ≤ 0.0001.

The finding that HDAC activity was elevated in *Rho*^I255d/+^ retina prompted us to test different HDAC inhibitors, targeting different HDAC classes and isoforms, for their potential to protect photoreceptors in the *Rho*^I255d/+^ ADRP model. We employed four different compounds, namely SAHA, MPT0G211 (MPT), ACY-957 (ACY), and NAM (Table 1), and first collected dose-response data in *Rho*^I255d/+^ retina using an *in situ* HDAC activity assay (Fig. S1A-H, Table S7). We found SAHA, MPT, and ACY treatment to generally decrease the numbers of ONL cells showing HDAC activity, however, exposure to NAM showed no clear effect and may even have increased net HDAC activity at intermediate compound concentrations. Having established the inhibitory capacities of these four compounds in mutant retina, we proceeded with evaluating their possible neuroprotective effects in organotypic retinal explant cultures.

### SAHA displays a narrow window-of-opportunity in *Rho*^I255d/+^ retinas

Previous studies found treatment with the HDAC class I and II inhibitor SAHA to delay photoreceptor cell death in ARRP models (14, 16). To assess the effect of SAHA on retinal degeneration in the *Rho*^I255d/+^ ADRP mouse, retinas were explanted at P12 and treated from P14 till P20 with either 0.01, 0.1, or 1 μM SAHA. WT retinal explants were included for comparison (24) (Table 1).

Cell death in ONL and inner nuclear layer (INL) was assessed using the TUNEL assay (Fig. 2A). NT mutant retina showed significantly more ONL TUNEL-positive cells than NT WT. At 0.1 μM, SAHA treatment decreased TUNEL-positive cell counts in *Rho*^I255d/+^ ONL (Fig. 2B, Table S2). However, at 1 μM, SAHA had toxic effects in WT ONL (Fig. 2B), even though there were no discernible changes in the INL in either WT or mutant retina (Fig. 2C, Table S2). NT mutant displayed a loss of ONL rows compared with NT WT (Fig. 2D, Table S2) and the ONL row count was reduced in both WT and *Rho*^I255d/+^ mutant retinas treated with 1 μM SAHA. These findings suggested that photoreceptor neuroprotection afforded by SAHA occurred only in a relatively narrow concentration window.

**Figure 2.**
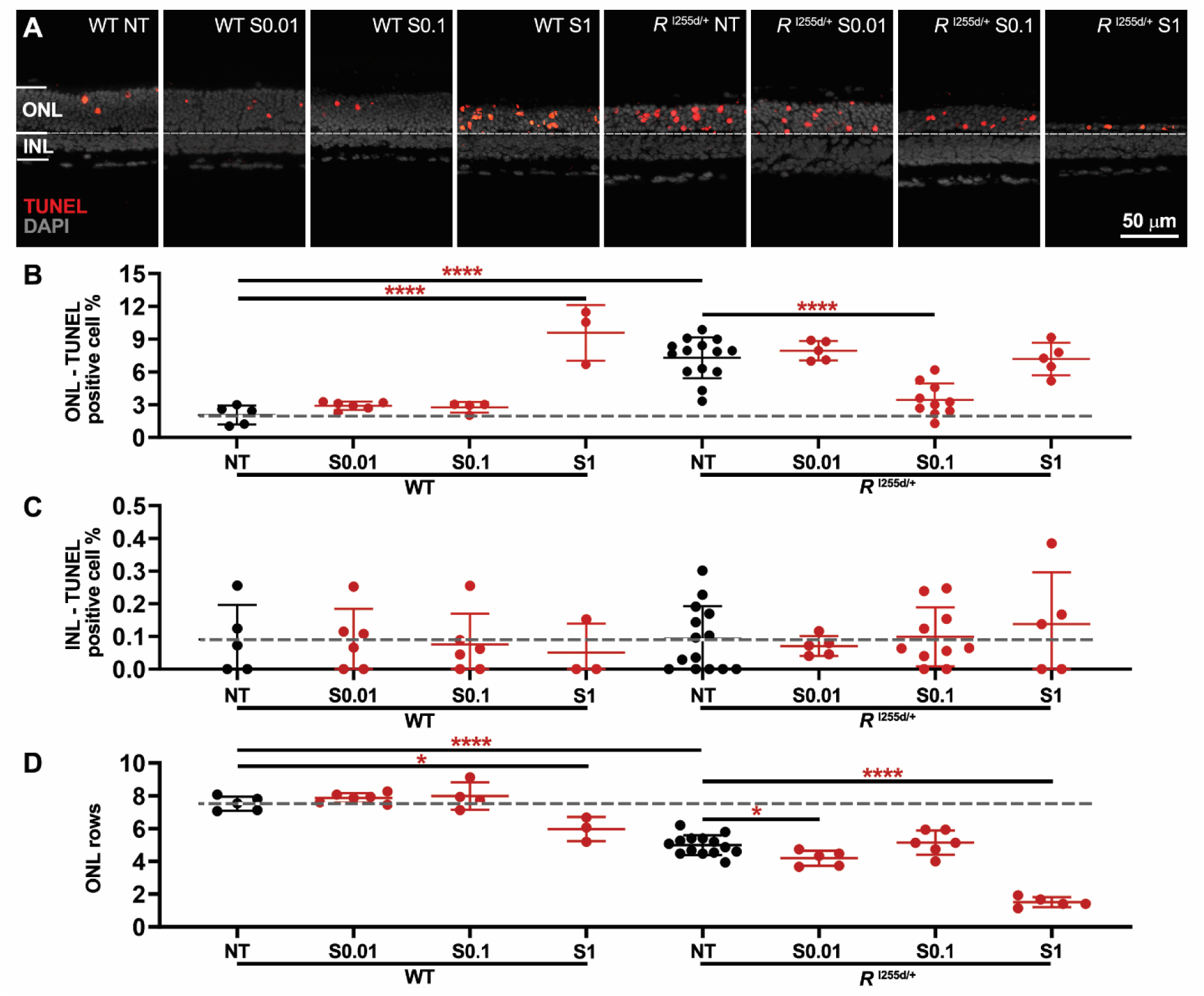
SAHA reduces *Rho*^I255d/+^ photoreceptor cell death in a narrow concentration range. Organotypic retinal explants were derived from wild-type (WT) and *Rho*^I255d/+^ (*R*^I255d/+^) mice and cultured from post-natal (P) day 12 to 20. Cultures were either non-treated (NT) or treated with different concentrations of SAHA (S; 0.01, 0.1, and 1 µM) from P14 onwards. **A.** Dying cells were visualized using the TUNEL assay (red) in the outer nuclear layer (ONL) and inner nuclear layer (INL) of both WT and *Rho*^I255d/+^ retina. DAPI (grey) was used as nuclear counterstain. **B**. In WT ONL, treatment with 1 µM SAHA increased TUNEL-positive cell counts, compared to NT WT. In *Rho*^I255d/+^ ONL, the numbers of dying cells were reduced at a SAHA concentration of 0.1 µM, compared to NT mutant. **C**. Percentage of TUNEL-positive cells in INL. SAHA treatment had no significant effects on the numbers of dying cells in WT and *Rho*^I255d/+^ INL. **D**. ONL row counts were decreased in NT *Rho*^I255d/+^ compared to WT, and were obviously reduced by 1 µM SAHA in both genotypes. Images represent results obtained from 3-14 independent retinal explant cultures; the data represent mean and SD; statistical analysis: Two-way ANOVA test with two-stage linear step-up procedure of Benjamini, Krieger, and Yekutieli; * = *p* ≤ 0.05, **** = *p* ≤ 0.0001.

### SAHA decreases apoptosis but increases non-apoptotic cell death, displays cone toxicity

Overactivation of HDACs may be linked to different neurodegenerative pathways (5, 6). In *Rho*^I255d/+^ retina, photoreceptor degeneration may be caused by both apoptosis and non-apoptotic cell death triggered by excessive cGMP-signaling (17, 18). A key event in apoptosis is the activation of the cysteine-type protease caspase-3 (5, 6), while activity of Ca^2+^-dependent calpain-2 has been connected to cGMP-dependent cell death (18, 25). To identify possible differential effects of SAHA treatment on apoptosis and cGMP-dependent cell death, we used immunostaining for activated caspase-3 and calpain-2 (Fig. 3, Table S3).

**Figure 3.**
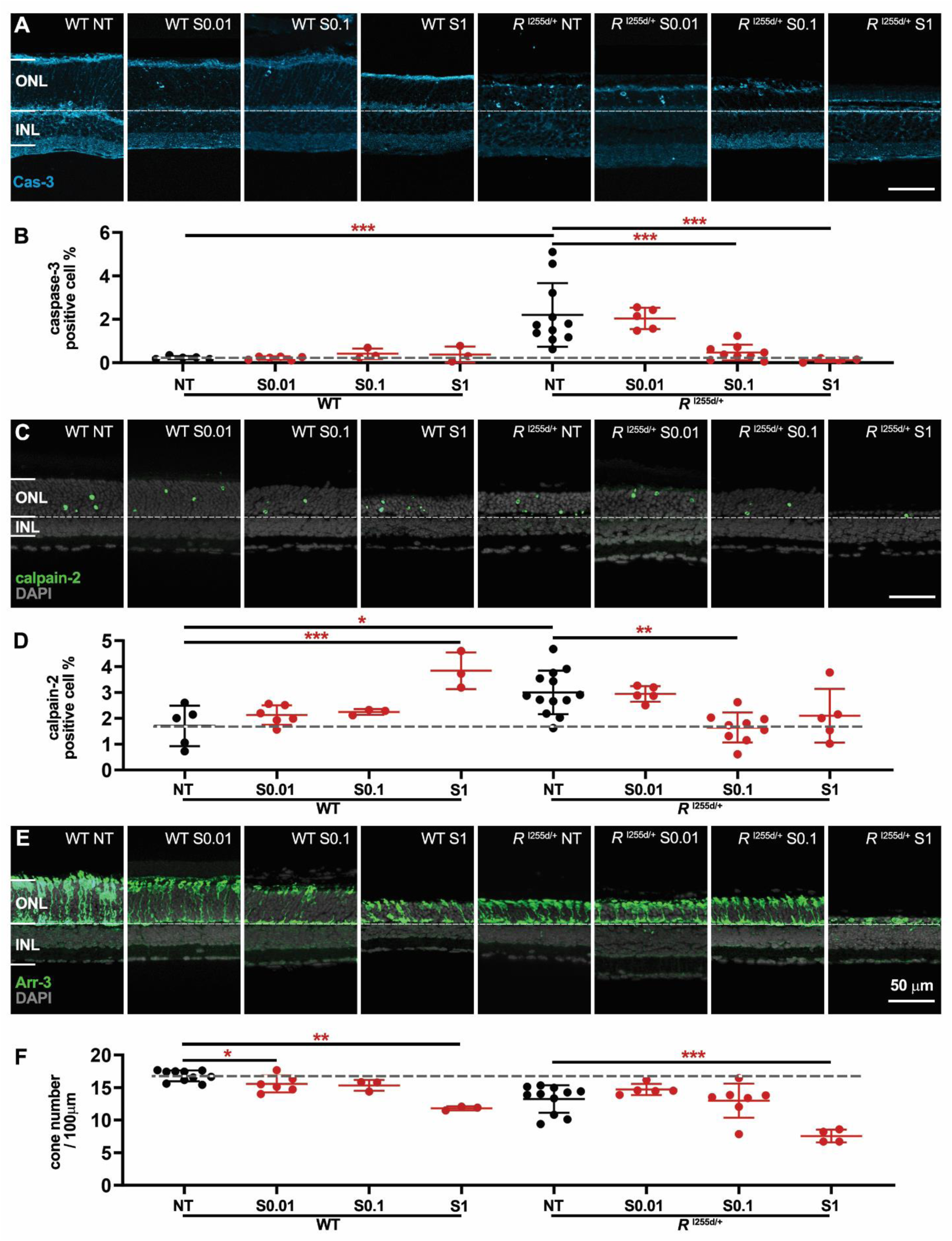
Differential effects of SAHA on markers for apoptotic and non-apoptotic cell death, cone toxicity. Organotypic retinal explants were derived from wild-type (WT) and *Rho*^I255d/+^ mice at post-natal (P) day 12 and cultured till P20. Retinal explants were treated with suberoyl-anilide hydroxamic acid (SAHA; S) in a concentration range from 0.01 to 1 µM. Immunostaining for cleaved caspase-3 (cyan), activated calpain-2 (green), and arrestin-3 (green) was used to detect cells undergoing apoptosis, non-apoptotic cell death, and cone photoreceptors, respectively. The non-treated (NT) WT situation served as reference (dashed lines in B, D, F). DAPI was used as nuclear counterstain (grey). **A, B.** SAHA treatment decreased the numbers of cells undergoing apoptosis in *Rho*^I255d/+^ outer nuclear layer (ONL) in a dose-dependent manner. **C, D**. The numbers of ONL cells displaying calpain-2 activation increased with SAHA concentration in WT retina. **E, F.** The numbers of cone photoreceptors per 100 µm of retinal circumference decreased with rising SAHA concentration in both WT and *Rho*^I255d/+^ mutant retina. Data represent mean and SD, and were obtained from 3-13 independent retinal explant cultures; statistical analysis: Two-way ANOVA test with two-stage linear step-up procedure of Benjamini, Krieger, and Yekutieli; * = *p* ≤ 0.05, ** = *p* ≤ 0.01, *** = *p* ≤ 0.001.

In organotypic retinal explants derived from WT and *Rho*^I255d/+^ mice at P12 and cultured until P20, the number of cells positive for activated caspase-3 was increased in NT mutant compared to NT WT, confirming earlier results (17) (Fig. 3A). SAHA treatment led to a dose-dependent decrease of caspase-3 positive cells in mutant ONL (Fig. 3B). This clear decrease in apoptosis contrasted with the rise in cell death as detected by the TUNEL assay (*cf*. Fig. 2), suggesting that SAHA triggered the execution of non-apoptotic cell death mechanisms.

Indeed, calpain-2 activation was increased in WT retinas treated with 1 μM SAHA (Fig. 3C, D), even though in mutant retinas at 0.1 μM SAHA, calpain-2 activation was decreased. Importantly, in both mutant *Rho*^I255d/+^ and WT retina, high levels of SAHA significantly reduced the number of arrestin-3 positive cone photoreceptors (Fig. 3E, F). Taken together, these results indicated that SAHA suppressed apoptosis but promoted non-apoptotic cell death, leading, notably, to a strong loss of cone photoreceptors.

### Rod and cone photoreceptors in *Rho*^I255d/+^ retina are protected by HDAC-6 inhibition

HDAC-6 plays a key part in cytoplasmic histone deacetylation (24, 26), however, its potential role in retinal degeneration remains unclear (27, 28). HDAC-6 is selectively inhibited by MPT0G211 (MPT; Table 1) (20, 24, 29). To investigate the impact of MPT treatment on *Rho*^I255d/+^ retinal degeneration, we proceeded as described above for SAHA, with concentrations of MPT ranging from 0.1 to 10 µM MPT (Fig. 4, Table S4; Fig. S2, Table S8).

**Figure 4.**
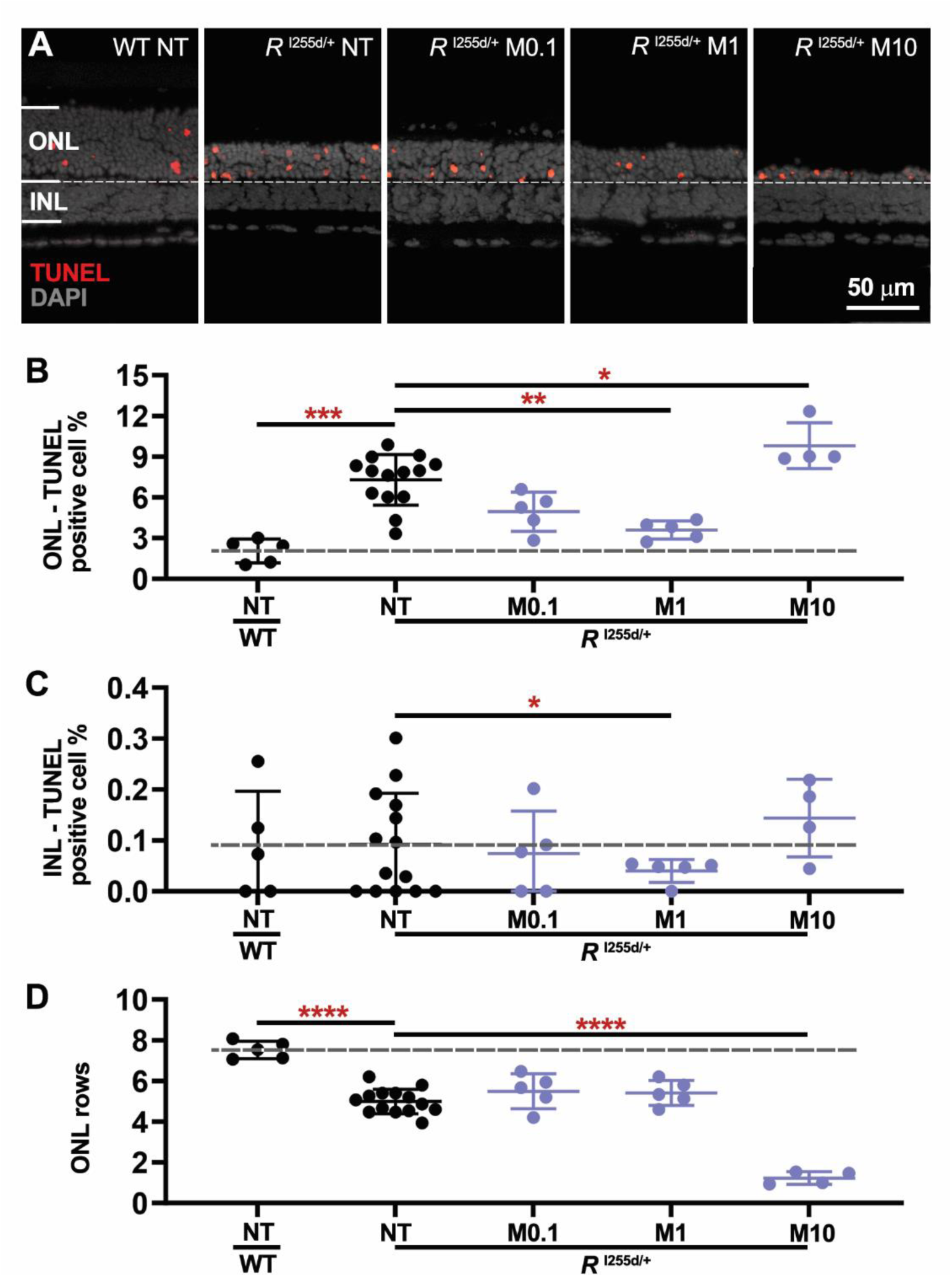
MPT reduces *Rho*^I255d/+^ cell death at low concentrations, but is cytotoxic at higher level. Organotypic retinal explants were derived from wild-type (WT) and *Rho*^I255d/+^ mice at post-natal (P) day 12 and cultured with MPT0G211 (MPT; M) from P14 to P20. The non-treated (NT) WT situation served as reference (dashed lines in B-D). **A.** The TUNEL assay (red) labeled dying cells in outer nuclear layer (ONL) and inner nuclear layer (INL) of both WT and *Rho*^I255d/+^ mutant retina. DAPI (grey) was used as nuclear counterstain. **B, C.** Quantification of TUNEL-positive cells in ONL (B) and INL (C). **D**. In *Rho*^I255d/+^ mutant retina, the ONL row count indicated significant photoreceptor loss at 10 µM MPT, when compared to NT mutant and WT. Data represent mean and SD, and were obtained from 4-14 independent retinal explant cultures; statistical analysis: Two-way ANOVA test with two-stage linear step-up procedure of Benjamini, Krieger, and Yekutieli; * = *p* ≤ 0.05, ** = *p* ≤ 0.01, *** = *p* ≤ 0.001, **** = *p* ≤ 0.0001.

In *Rho*^I255d/+^ mutant ONL, the number of dying cells were reduced by treatment with 1 µM MPT (Fig. 4A, B). However, ONL cell death was significantly increased when mutant retina was exposed to 10 µM MPT. INL cell death was reduced by 1 µM MPT treatment (Fig. 4C), while the number of ONL cell rows was strongly decreased by 10 µM MPT (Fig. 4D).

At 10 µM concentration, MPT increased markers for both apoptotic and non-apoptotic cell death (Fig. S2A-D). Remarkably, at low to intermediate concentrations of 0.1 and 1 µM, MPT treatment significantly preserved cone photoreceptors, an effect that was reversed at 10 μM concentration (Fig. S2E, F). These findings suggested that from 0.1 – 1 µM MPT treatment afforded neuroprotection of both rod and cone photoreceptors, but that at higher levels, MPT was toxic to *Rho*^I255d/+^ retina. Moreover, HDAC-6 activity apparently contributed to *Rho*^I255d/+^ mutation-induced neurodegeneration.

### ACY protects *Rho*^I255d/+^ cone photoreceptors, switches non-apoptotic cell death to apoptosis

HDAC-1 and -2 are regarded as the major isoforms regulating nuclear deacetylation (24). To investigate the role of HDAC-1 and -2 in *Rho*^I255d/+^ retina, we employed the highly specific inhibitor ACY-957 (ACY, Table 1) (24, 30) at concentrations between 0.01 and 10 μM (Fig. 5, Table S5; Fig. S3, Table S9).

**Figure 5.**
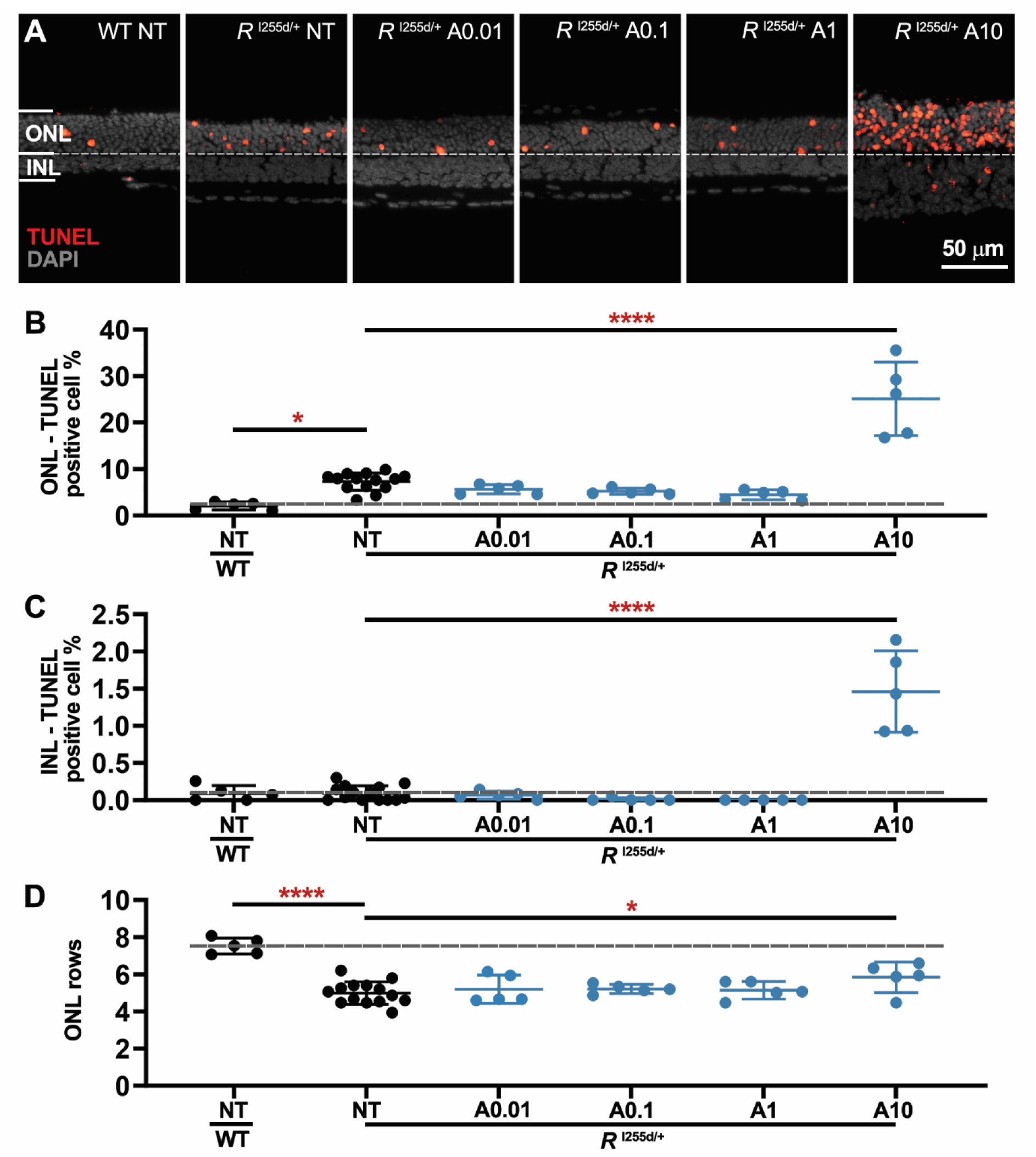
ACY is cytotoxic to *Rho*^I255d/+^ retina at high concentration. Organotypic retinal explants were derived from wild-type (WT) and *Rho*^I255d/+^ mice at post-natal (P) day 12 and cultured with ACY-957 (ACY; A) from P14 to P20. The non-treated (NT) WT situation was used as reference (dashed lines in B-D). **A.** The TUNEL assay (red) labeled dying cells in outer nuclear layer (ONL) and inner nuclear layer (INL) of both WT and *Rho*^I255d/+^ mutant retina. DAPI (grey) was used as nuclear counterstain. **B, C.** Quantification of TUNEL-positive cells in ONL (B) and INL (C). Note the strong rise in TUNEL positive cells at 10 µM ACY concentration. **D**. In *Rho*^I255d/+^ mutant retina, the ONL row count indicated a significant preservation of photoreceptors at 10 µM ACY, when compared to NT mutant. Data represent mean and SD, and were obtained from 5-14 independent retinal explant cultures; statistical analysis: Two-way ANOVA test with two-stage linear step-up procedure of Benjamini, Krieger, and Yekutieli; * = *p* ≤ 0.05, ****= *p* ≤ 0.0001.

TUNEL assay results (Fig. 5A-C) showed no major effects on ONL and INL cell death at ACY concentrations from 0.01 to 1 µM. However, there was a marked increase in ONL and INL cell death at 10 µM. Paradoxically, at 10 µM there was also an increase in ONL row counts (Fig. 5D), *i.e.* a finding that seemingly contradicted the high TUNEL positive cell counts.

Moreover, at relatively low concentrations of 0.01 μM and 0.1 μM ACY treatment decreased the numbers of caspase-3 activation positive cells in the *Rho*^I255d/+^ mutant ONL to essentially zero, indicating a complete suppression of apoptosis (Fig. S3A, B). Remarkably, at 10 µM, ACY treatment strongly increased caspase-3 activation, concomitant with a decrease in calpain-2 activation (Fig. S3C, D). Finally, ACY significantly preserved cones at 0.1 and 1 µM concentration (Fig. S3E, F).

### Nicotinamide treatment causes selective rod photoreceptor toxicity

Nicotinamide (NAM) specifically inhibits NAD^+^-dependent HDACs belonging to class III, *i.e.* sirtuin-type HDACs (Table 1) (22, 23). To study the effects of NAM on *Rho*^I255d/+^ retina, we proceeded as described above for SAHA, with NAM concentrations between 20 to 2000 µM (Fig. 6, Table S6; Fig. S4, Table S10).

**Figure 6.**
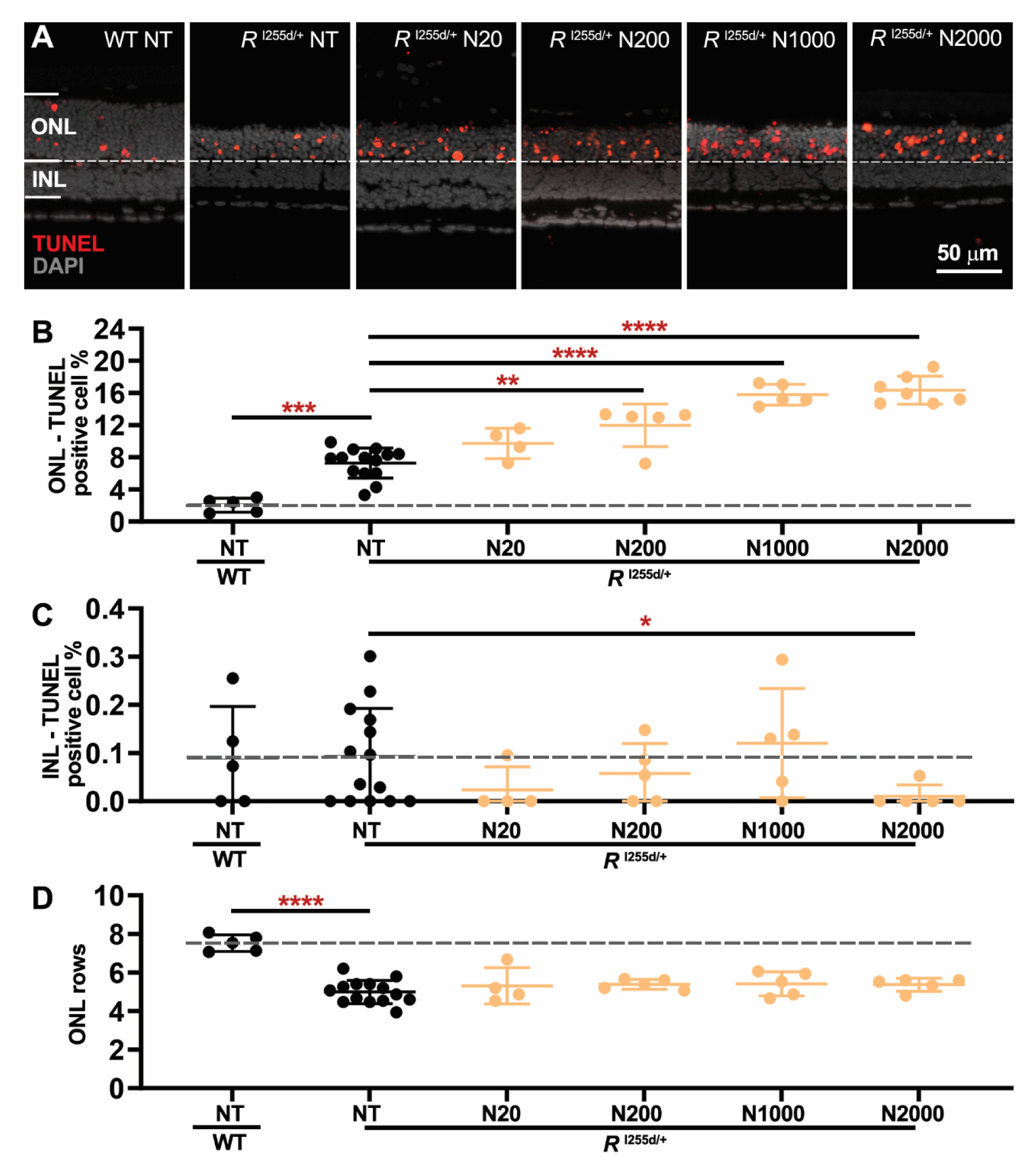
NAM induces photoreceptor cell death in *Rho*^I255d/+^ retina. Organotypic retinal explants were derived from wild-type (WT) and *Rho*^I255d/+^ mice at post-natal (P) day 12 and treated with 20 to 2000 µM nicotinamide (NAM, N) from P14 to P20. The non-treated (NT) WT situation was used as reference (dashed lines in B-D). **A.** The TUNEL assay (red) labeled dying cells in outer nuclear layer (ONL) and inner nuclear layer (INL) of both WT and *Rho*^I255d/+^ mutant retina. DAPI (grey) was used as nuclear counterstain. **B, C.** Quantification of TUNEL-positive cells in ONL (B) and INL (C). Note the strong rise in TUNEL positive cells seen in ONL but not in INL. **D**. In *Rho*^I255d/+^ mutant retina, the ONL row count with NAM treatment remained at the level of NT mutant. Data represent mean and SD, and were obtained from 5-14 independent retinal explant cultures; statistical analysis: Two-way ANOVA with two-stage linear step-up procedure of Benjamini, Krieger, and Yekutieli; * = *p* ≤ 0.05, ** = = *p* ≤ 0.01, *** = *p* ≤ 0.001, **** = *p* ≤ 0.0001.

TUNEL staining showed that NAM treatment promoted cell death, particularly in *Rho*^I255d/+^ ONL (Fig. 6A-B). In contrast, INL neurons were not affected by NAM, and there was even a slight decrease in the number of dying cells at 2000 µM (Fig. 6C). Under NAM treatment the ONL row count remained unchanged compared to NT *Rho*^I255d/+^ explants (Fig. 6D).

While caspase-3 activation was increased at 2000 μM NAM, calpain-2 activation was not changed in *Rho*^I255d/+^ retina at any NAM concentration employed (Fig. S4A-D). Also, the number of cone photoreceptors was not changed by NAM treatment (Fig. S4E, F). Together, these findings suggested that *Rho*^I255d/+^ rod photoreceptors – but no other retinal cell type – were especially vulnerable to NAM, and, by inference, dependent on NAD^+^-related processes.

### ACY can promote ONL cell proliferation

Numerous studies have connected HDAC activity to cell death (24), yet HDACs may also govern cell proliferation (31). To assess possible pro-proliferative effects of HDAC inhibition, we exposed inhibitor-treated *Rho*^I255d/+^ retinas to 10 μM bromodeoxyuridine (BrdU) from P16 to P18 (32–34). The incorporation of BrdU into the DNA of dividing cells was later visualized using anti-BrdU immunostaining.

Treatment with MPT and NAM did not lead to appreciable BrdU labeling in *Rho*^I255d/+^ retina, yet SAHA caused a minor (non-significant) increase in ONL BrdU positive cells (Fig. S5A, B, Table S11). In contrast, ACY treatment significantly increased BrdU-positive cells in *Rho*^I255d/+^ ONL. This outcome may explain the seemingly contradictory results where ACY treatment caused a large increase in ONL cell death, as evidenced by the TUNEL assay, concomitant with an increase in ONL cell rows (*cf*. Fig. 5). The cell division shown by BrdU-labeling is remarkable for a neuronal tissue that is generally considered to be post-mitotic, and suggests that some retinal cells may become proliferative when HDAC-1 and/or HDAC-2 are inhibited.

### Overview of HDAC inhibition effects in *Rho*^I255d/+^ retina

The compounds SAHA, MPT, and ACY selectively inhibit class I/II/IV HDAC isoforms at low nanomolar concentrations (Table 1). To facilitate a comparative analysis, dose-response curves for the treatment effects on ONL cell death, ONL row counts, cone numbers, HDAC activity, as well as activation of caspase-3 and calpain-2 are shown in Figure 7. At low to intermediate concentrations, treatment with SAHA and MPT decreased the numbers of TUNEL-positive cells in *Rho*^I255d/+^ ONL (Fig. 7A). Increased ONL row counts were observed only with high-dose ACY treatment, perhaps due to increased cell proliferation (Fig. 7B). A significant cone preservation was observed at intermediate doses of MPT and ACY (Fig. 7C). SAHA, MPT, and ACY all decreased HDAC activity positive cells in the ONL, with the pan-inhibitor SAHA showing the strongest, and the HDAC-6 specific inhibitor MPT the weakest overall effect (Fig. 7D). At intermediate concentrations, both SAHA and ACY strongly reduced caspase-3 activation and hence apoptosis (Fig. 7E). The activation of calpain-2 was decreased by intermediate SAHA levels and by high concentration ACY (Fig. 7F). Overall, in terms of photoreceptor neuroprotection, the HDAC-6 inhibitor MPT displayed the most pronounced effects, with significant reduction of ONL cell death and improved cone survival in the 0.1 to 1 µM concentration range.

**Figure 7.**
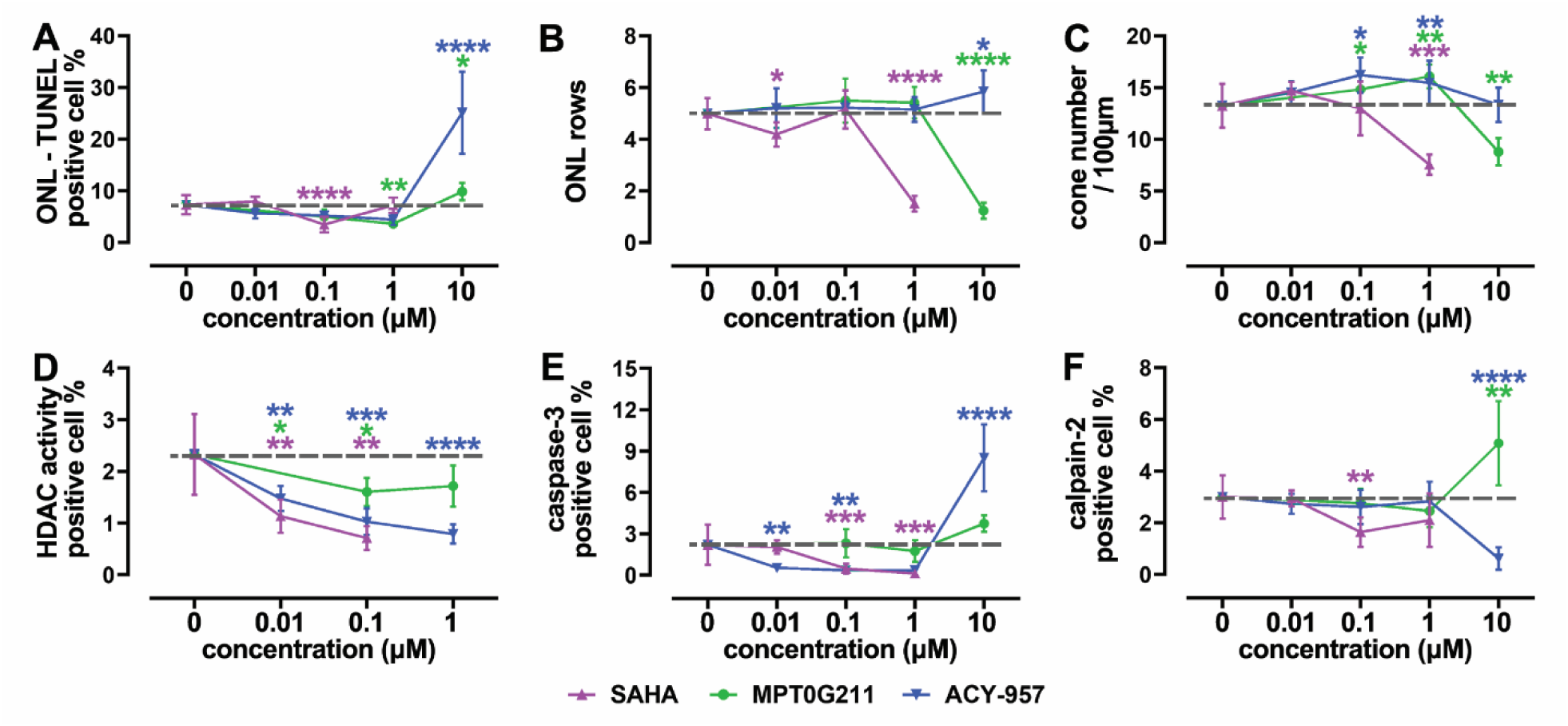
Overview of HDAC inhibition effects in *Rho*^I255d/+^ retina. Dose-response curves (log scale) for SAHA (magenta), MPT0G211 (green), and ACY-957 (blue). **A**. Effects on cell death as evidenced by the TUNEL assay. **B**. Photoreceptor survival as assessed by outer nuclear layer (ONL) row counts. **C**. Numbers of cone photoreceptors per 100 µM of retinal circumference. **D**. Quantification of HDAC activity positive cells in the ONL. **E**. Quantification of ONL cells showing caspase-3 activation. **F.** Quantification of ONL cells displaying calpain-2 activation. Data obtained from 5-13 independent retinal explant cultures; data points indicate mean and SD; statistical analysis: Two-way ANOVA test with two-stage linear step-up procedure of Benjamini, Krieger, and Yekutieli; * = *p* ≤ 0.05; ** = *p* ≤ 0.01; *** = *p* ≤ 0.001, **** = *p* ≤ 0.0001.

## Discussion

Aberrant HDAC activity has been linked to photoreceptor degeneration, mostly in models for ARRP, and HDAC inhibitors were suggested as possible therapeutic agents for IRD therapy (5). Here, we show that increased HDAC activity also occurs during photoreceptor death in the human homologous *Rho*^I255d/+^ ADRP mouse model. We found HDAC inhibition to have both protective and destructive effects, with the HDAC-6 inhibitor MPT showing the most promising results overall.

### HDAC inhibition differentially affects *Rho*^I255d^**^/+^** rod and cone photoreceptors

Earlier studies have shown HDAC activity to be involved in photoreceptor cell death (5, 6), including in animal models for ARRP (*e.g*., *rd1*, *rd10* mice) and ADRP (*e.g*., *Rho*^S334ter^, *Rho*^P23H^ rats) (5). However, the specific roles of individual HDAC isoforms remain elusive. HDAC-1, -2, and -3 belong to class I and primarily deacetylate nuclear proteins, while the class II HDAC-6 is responsible for deacetylation in the cytoplasm (24, 35). Increased HDAC activity in the *Rho*^I255d/+^ retina primarily involved class I and II isoforms, aligning with previous results obtained in *rd1* retina and indicating causal involvement of class I and II isoforms in both ARRP and ADRP. Inhibition of class I and II HDACs by TSA or SAHA was shown to promote photoreceptor viability in various IRD models. This related to both cone protection in the *cpfl1* mouse and rod protection in *rd1* and *rd10* mice (11–16). In the present study on *Rho*^I255d/+^ retina, SAHA protected mainly rods (albeit only in a narrow concentration range), while HDAC-1 and -2 inhibition by ACY primarily delayed cone degeneration. Importantly, the HDAC-6 specific inhibitor MPT preserved both rods and cones, confirming an earlier *rd10* mouse *in vivo* study, which found HDAC-6 inhibition to delay mostly cone degeneration (36).

Altogether, specific inhibition of HDAC isoforms -1, -2, and -6 may develop into a mutation-independent therapeutic strategy applicable to both ARRP and ADRP. Remarkably, none of the HDAC inhibitors tested affected INL neurons (except ACY at high concentration), indicating that photoreceptor physiology was more sensitive to changes in HDAC activity, a phenomenon perhaps related to the very high metabolic rates of photoreceptors (37, 38).

### Caspase-3 *vs*. calpain-2: Apoptosis *vs*. non-apoptotic cell death

Different HDAC isoforms may play important roles in apoptosis or in non-apoptotic cell death triggered, for instance, by excessive cGMP-signaling. Increased activation of caspase-3 as a marker for apoptosis was found in dying photoreceptors in the *Rho*^S334ter^ rat (5) and the *Rho*^I255d/+^ mouse models for ADRP (17). Caspase-3 can specifically cleave certain HDAC isoforms, including HDAC-3 and HDAC-4, which may further promote apoptosis (39, 40). Still, in many forms of retinal degeneration caspase-3 activation, and the execution of apoptosis overall, does not seem to play a major role (5, 41). This is confirmed by our data on the effects of SAHA and ACY on caspase-3: Both compounds robustly reduced caspase-3 activity, yet they were unable to prevent mutation-induced photoreceptor death.

Apart from caspase-3, also activity of Ca^2+^-dependent calpain-type proteases was linked to retinal degeneration, in many different ARRP and ADRP animal models, and concomitant with increased HDAC activity (5). In relative terms, the numbers of dying cells corresponded well to the numbers of cells exhibiting activation of calpain-2. Calpain-2 activation is regarded as a relatively short-lived event occurring towards the end of the cell death process (42) and is recognized as a neurodegenerative factor in ARRP models (25, 43), linked to non-apoptotic forms of cell death (44). Curiously, *Rho*^I255d/+^ photoreceptors displayed significant activation of both caspase-3 and calpain-2, implying parallel execution of apoptotic and non-apoptotic mechanisms (17, 18). Since the simultaneous execution of apoptotic and non-apoptotic cell death would be metabolically incompatible (6), this indicates that different populations of photoreceptors use different mechanisms, perhaps depending on the momentary metabolic status at the time cell death was triggered. At any rate, our data suggests that HDAC inhibitors can shift the balance between apoptosis and non-apoptotic cell death. For instance, activation of caspase-3 was decreased by the pan-inhibitor SAHA and the HDAC-1/-2 inhibitor ACY at low concentrations, but increased by ACY at high concentration. Thus, in *Rho*^I255d/+^ retina HDAC-1 and -2 may promote photoreceptor apoptosis. Then again, high concentration ACY treatment strongly increased caspase-3 but decreased calpain-2 activation. This switch from non-apoptotic to apoptotic cell death could be dependent on HDAC-3, inhibited by ACY only at high concentrations.

### NAM and NAD^+^: Impact on metabolism and cell death

NAM is a non-competitive inhibitor of sirtuin-type, class III HDAC isoforms where it binds and blocks the active site (45). NAM can induce oxidative stress and damage mitochondria, leading to loss of mitochondrial membrane potential and cell death (46). Given their high energy demand (37), it is easy to see why photoreceptors would be more affected by NAM treatment than other retinal cell types. In *Rho*^I255d/+^ retina, cell death induced by NAM treatment was restricted to rod photoreceptors, suggesting important roles for sirtuins in rods. This result may also be indicative of significant differences in the metabolism of rods and cones (47, 48), and could relate to the mitochondrial sirtuin isoforms -3, -4, and -5, thought to regulate key metabolic tasks, such as antioxidant defense and energy production (49). In a recent study, rod energy production was found to be reliant on mitochondrial oxidative phosphorylation, while cones were able to survive using only glycolysis (47, 50). It is thus plausible to think that a blockage of mitochondrial sirtuins could have strong detrimental effects on rods but leave cones unaffected.

Since NAM also inhibits poly(ADP-ribose)polymerase (PARP) with similar efficacy as sirtuins (*cf*. Table 1), an alternative explanation should also be considered: The PARP family of enzymes is involved in regulating gene transcription and promoting DNA damage repair (51). NAM competitively inhibits PARP, and the resulting lack of gene transcription and DNA repair may be especially detrimental for photoreceptor cells with their very high transcriptional activity. The most transcribed gene in rods (and possibly the most transcribed gene in any cell type) is *Rho* (52) and this could make the *Rho*^I255d^ mutant studied here particularly vulnerable to PARP inhibition.

### Role of HDACs in cell proliferation

HDAC-1 is known to play a critical role in cell proliferation since its absence can cause cell cycle arrest, reduce numbers of mitotic cells, and inhibit cell growth (53). Accordingly, HDAC inhibition can have antiproliferative effects, primarily by inducing cell-cycle arrest (54–56), which in a post-mitotic neuronal tissue such as the retina should not be problematic.

BrdU labeling showed that neither SAHA, MPT, nor NAM induced cell proliferation in *Rho*^I255d/+^ retina. However, high doses of ACY resulted in increased ONL cell proliferation, as indicated by elevated numbers of BrdU-positive cells and confirmed by higher ONL row counts. Since ACY targets HDAC-1 and HDAC-2 rather selectively but can also block HDAC-3 at ≈100-fold higher concentrations (Table 1), this outcome could indicate anti- and pro-proliferative functions of HDAC-1/HDAC-2 and HDAC-3, respectively.

Paradoxically, high dose ACY also induced caspase-3 positive apoptosis in *Rho*^I255d/+^ ONL. This may correspond to similar findings in HDAC-1 mutant zebrafish, where increased retinal apoptosis was associated with proliferation, indicating that HDAC-1 is cell-autonomously required for retinal cell-cycle exit and differentiation (57). Alternatively, non-cell-autonomous cell proliferation of Müller glia cells may have been induced by cell death triggered by high-dose ACY treatment (58). Independent of this, our BrdU data may have ramifications for therapies that aim at replacing lost cells by stimulating endogenous regeneration (59).

## Conclusion

Our study illustrates the complexity of HDAC functions in photoreceptor cell death and survival. While HDAC inhibitors can potentially preserve photoreceptors in the *Rho*^I255d/+^ mouse model for ADRP, their effects are ambiguous and need to be carefully evaluated for individual HDAC isoforms. Even though the pan-inhibitor SAHA exhibited robust anti-apoptotic properties, its net treatment effects were detrimental. The HDAC-1 and -2 inhibitor ACY displayed relatively strong cone protection, and MPT, targeting HDAC-6, preserved both rod and cone photoreceptors. Our data furthermore suggest that rods and cones diverge in terms of NAD^+^ consumption. In summary, HDAC inhibitors present a promising avenue for future therapeutic development, both for ARRP and ADRP. Future research may further narrow-down disease-relevant HDAC isoforms and elucidate underlying degenerative mechanisms to optimize therapeutic efficacy.

## Materials and Methods

### Animals

In the present study, we used C57BL/6J wild-type (WT) and heterozygous *Rho*^I255d/+^ (*R*^I255d/+^) mice for this project (17). Animals were housed under standard white cyclic lighting, had free access to food and water, and were used regardless of gender. All efforts were made to minimize the number of animals used and their suffering. The total number of mice (retinal explants) used for this study was 86 (152).

### *In vitro* organotypic retinal explant culture procedure

To assess the effects of HDAC inhibitors, organotypic retinal explants were prepared as previously described (60). Briefly, *Rho*^I255d/+^ and WT mice were sacrificed at post-natal (P) day 12 through CO_2_ inhalation. Their eyeballs were placed in R16 basal medium (BM, 074-91252A, Gibco, Paisley, UK) for 5 min at room temperature (RT), and later incubated in R16 BM medium with 0.12% proteinase K (39450, MP Biomedicals, Irvine, CA, USA) at 37 °C for 17 min. Proteinase K was then inactivated by adding 20% fetal bovine serum (FBS, F7524, Sigma, Hamburg, Germany) in BM at RT. The sclera and anterior segments were carefully removed from the eyeballs, and the optic nerve was cut, leaving the RPE attached to the retinas. Retinas were cut into a four-leaf clover shape and transferred to a culture membrane insert in a six-well plate (Corning Life Sciences, Corning, NY, USA) with RPE facing down. Retinal explants were incubated in 1.2 mL of R16 complete medium (CM) with supplements, under completely defined conditions, free of serum and antibiotics.

### HDAC inhibitor treatment

Explants were incubated in a sterilized and humidified incubator with 5% CO_2_ at 37 °C. After 48h, the retinas were either kept as non-treated (NT) control, or exposed to the HDAC inhibitors SAHA (0.01, 0.1, and 1 μM), MPT0G211 (MPT; 0.1, 1, and 10 μM), ACY-957 (ACY; 0.01, 0.1, 1, and 10 μM), or nicotinamide (NAM; 20, 200, 1000 and 2000 μM) (Table 1; dissolved in dimethyl sulfoxide (DMSO)). DMSO was found to be non-toxic to the retina for concentrations of up to 1% (unpublished observations). To address possible cellular proliferation induced by HDAC inhibition, we incubated retinal explants with an additional 10 μM 5-bromo-2′-deoxyuridine (BrdU) for two days, from P16 to P18. This was followed by BrdU-immunolabelling (see below). The medium was changed every second day, and cultures ended at P20.

Mouse retinal explants derived from the left and right eyes of a given animal were separated into two groups and consecutively numbered. The assignment to either the NT or experimental group was determined by rolling a die for the left and right eyes of each animal. However, the allocation was consistently arranged so that the two retinas from the same animal were always assigned to different groups. No blinding procedures were implemented for sample allocation.

### Histology

#### Fixed sections

For retinal cross-section preparation, eyeballs were obtained from P20 *Rho*^I255d/+^ and WT mice. A perforation was made from the limbus toward the vitreous cavity and fixed in 4% paraformaldehyde (PFA) for 45 min. Afterwards, the anterior segment, lens, and vitreous body were removed. Retinal explant cultures were terminated at P20 and then fixed directly in 4% PFA for 45 minutes. Both eyecups and retinal explants were cryoprotected with graded sucrose solutions (10%, 20%, and 30%), and later on, embedded in Tissue-Tek O.C.T. compound (Sakura Finetek Europe, Alphen aan den Rijn, Netherlands). The retina blocks were cryosectioned sagittally with 12 µm thickness and prepared on a Thermo Scientific NX50 microtome (Thermo Scientific, Waltham, MA), then collected on Superfrost Plus glass slides (R. Langenbrinck, Emmendingen, Germany), then stored at −20 °C.

### TUNEL assay

Photoreceptor cell death was examined utilizing a terminal deoxynucleotidyl transferase dUTP nick end labeling (TUNEL) assay (Sigma-Aldrich *In Situ* Cell Death Detection Kit, 11684795910, red fluorescence)(61). Fixed retinal tissue sections were dried at 37°C for 30 min, then rinsed with phosphate-buffered saline (PBS) solution at RT for 10 min. Sections were subsequently incubated in Tris-buffered saline (TBS) containing proteinase K at 37°C for 5 min to inactivate nucleases. After 3 washes in TBS for 5 min each, sections were placed in an ethanol-acetic acid mixture (70:30) at -20°C for 5 min, followed by 3 additional washes in TBS. They were then incubated in blocking solution (comprising 10% normal goat serum, 1% bovine serum albumin, and 1% fish gelatin in 0.1% PBS-Triton X-100) for 1 h at RT. Subsequently, sections were stained using the kit as per the manufacturer’s instructions and incubated in assay solution overnight at 4°C. The following day, sections were washed 2 times in PBS for 5min each. Finally, tissue sections were mounted with Vectashield (Vector Laboratories, Newark, CA, USA), containing 4′,6-diamidino-2-phenylindole (DAPI) as nuclear counterstain. Stained sections were imaged under a Zeiss (ApoTome 2) microscope.

### HDAC activity

The HDAC activity assay was adapted from the FLUOR DE LYS®-Green System (Biomol, Hamburg, Germany) and performed on 4% PFA-fixed cryosections from both *Rho*^I255d/+^ and WT animals. Sections were dried at 37°C for 30 min and then placed in HDAC assay buffer (50 mM Tris-HCl, 137 mM NaCl, 2.7 mM KCl, 1 mM MgCl2, and 10% bovine serum albumin (BSA)) for 30 min. Next, they were exposed to the fluor-de-lys Sirt-2 substrate working solution (for 100 µL: 3 µL FLUOR DE LYS®-SIRT2 deacetylase substrate (200 µM; Enzo Life Sciences, New York, USA) + 4 µL NAD+ (1mM) + 1 µL NP-40 (0.1%) + 92 µL assay buffer) for 3 h at 37°C. After washing once with PBS for 5 min, the slides were fixed in methanol for 20 min at -20°C. After being washed once with PBS for 5 min, the slides were stained with ToPro3 (1:1000, T3605, Invitrogen, Carlsbad, USA) for 20 min. Afterward, the slides were washed with PBS and mounted with FLUOR DE LYS developer II concentrate (BML-KI176-1250, Enzo Life Sciences, New York, USA), incubated overnight and imaged the next day. To investigate the activity of HDAC *ex vivo* (15), cryosections of eyecups were additionally incubated with HDAC inhibitors (Table 1; 2 μM Trichostatin A; TSA, 2 mM nicotinamide; NAM, and 2 μM TSA + 2 mM NAM) in the incubation of HDAC assay buffer, working solution, and developer II. Negative controls were made by omitting the HDAC substrate.

### Immunofluorescence

The fixed cryosections were dried for 30 min at 37°C, washed with PBS 1 time for 10 min, incubated in a blocking solution (10% NGS and 1% BSA in PBS with 0.3% Triton X-100 (0.3% PBST)) for 1 h at RT. Primary antibodies used were: rabbit anti-cleaved caspase-3 (1:100, 5A1E, cell signaling, Danvers, USA), rabbit anti-calpain-2 (1:200, ab39165, Abcam, Cambridge, UK), rabbit anti-cone arrestin-3 (1:500, ab15282, EMD, Millipore Corp., USA), and rabbit anti-BrdU (1:100, ab6326, Abcam, Cambridge, UK) were diluted in blocking solution and incubated with the sections overnight at 4°C. The secondary antibody (1:350, goat anti-rabbit Alexa Fluor 488, Molecular Probes, Eugene, OR) was incubated for 1 h, then washed 3 times in PBS and mounted with Vectashield with DAPI (Vector Laboratories, Newark, CA, USA), stored at 4 °C before imaging.

### Microscopy

The images of sections were captured using a Zeiss Imager Z.2 fluorescence microscope equipped with ApoTome2, an Axiocam 506 mono camera, and HXP-120V fluorescent lamp (Carl Zeiss Microscopy, Oberkochen, Germany). The excitation (λExc.)/emission (λEm.) characteristics of the filter sets used for the different fluorophores were as follows (in nm): DAPI (λExc. = 369 nm, λEm = 465 nm), AF488 (λExc. = 490 nm, λEm = 525 nm), and ToPro3 (λExc. = 642 nm, λEm = 661 nm). Images were captured from the retina using a 20x/0.8 Zen 2.3 blue edition microscope and software. Sections of 12µm thickness were analyzed using 9 ApoTome2 Z-stack. Final figures for publication were processed and assembled by Adobe Photoshop 2020 (San Jose, CA, USA).

### Quantification

For each datapoint and staining procedure (HDAC activity, TUNEL assay, cleaved caspase-3, calpain-2, cone arrestin-3, and BrdU), images were captured on 3 retinal sections, at positions between +/- 30° lateral to the optic nerve from at least 3 different animals per genotype. The average area occupied by a photoreceptor cell (*i.e.,* cell size) was determined by counting DAPI- or ToPro3-stained nuclei in 9 different regions of the retinal outer nuclear layer (ONL) to calculate the average cell size. The percentage of positive cells was calculated by the absolute number of positive cells divided by the total number of ONL cells. The quantification of cones was presented as the number of cones per 100 μm of retinal circumference.

### Statistics

To determine the minimum number of animals and samples required for the statistical analyses, a power calculation based on previously observed effect sizes (25, 62) was performed using the online tool StatistikGuru, Version 1.96 (www.statistikguru.de; information retrieved July 2025). Animals were not assigned to experimental groups prior to their sacrifice, both retinal explants generated from experimental animals were included in the analysis. Errors in graphs and text are given as standard deviation (SD). Values are given as mean ± SD. Statistical analysis was performed using GraphPad Prism 10.1.2 software (GraphPad Software, La Jolla, CA, USA), Two-way ANOVA test with the two-stage linear step-up procedure of Benjamini, Krieger and Yekutieli was performed to compare more than two groups. Levels of significance were: * = *p* ≤ 0.05, ** = *p* ≤ 0.01, *** = *p* ≤ 0.001, **** = *p* ≤ 0.0001. Full statistical reports are presented in Tables S1-11.

## Acknowledgments

The authors would like to thank Norman Rieger (Institute for Ophthalmic Research, Eberhard-Karls-Universität Tübingen) for excellent technical assistance.

## Author contributions

Conceptualization, Y.Z. and F.P.-D.; methodology, Y.Z., and P.N.-K.; visualization, Y.Z.; software, Y.Z.; validation, Y.Z.; formal analysis, Y.Z.; investigation, Y.Z. and P.N.-K.; data curation, Y.Z.; writing-original draft preparation, Y.Z.; writing—review and editing, K.W.J. and F.P.-D.; supervision, F.P.-D.; project administration, F.P.-D.; funding acquisition, F.P.-D. All authors have read and agreed to the published version of the manuscript.

## Data Availability Statement

All data generated or analyzed during this study are included in this published article and its Supplementary Materials Files.

## Ethics approval and consent to participate

All procedures were performed using the ARVO statement for using animals in ophthalmic and visual research. Animal protocols compliant with §4 of the German law of animal protection were reviewed and approved by the Tübingen University committee on animal protection (Einrichtung für Tierschutz, Tierärztlicher Dienst und Labortierkunde, Registration No. AK02/19M, AK01/22M).

