## supplemental figures for "Differential effects of HDAC inhibitors in the *Rho*^I255d^ mouse model for autosomal dominant retinitis pigmentosa"

#### Figure S1

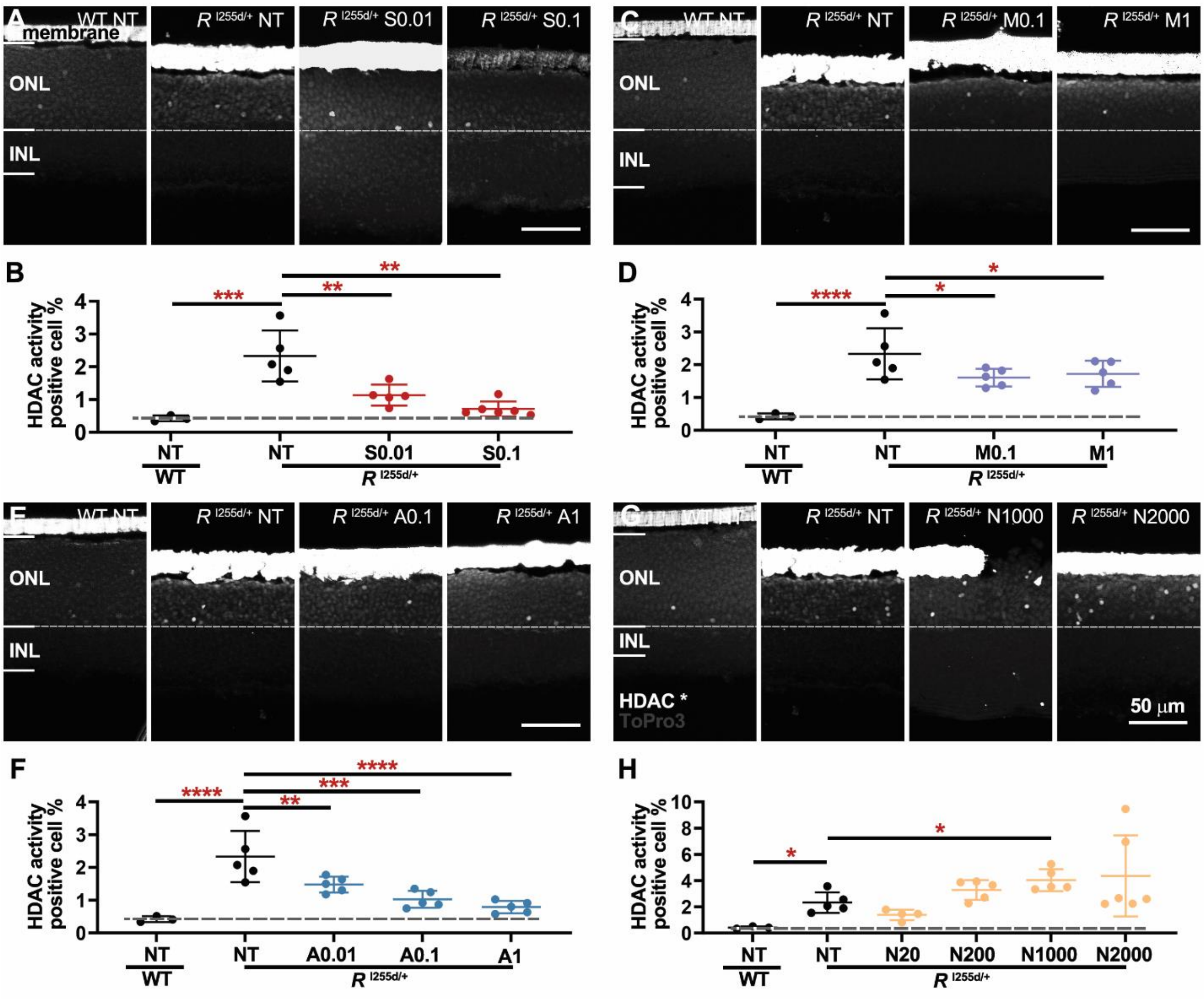


**Figure S1. In *Rho*$^{I255d/+}$ retina HDAC activity was decreased by SAHA and ACY, but increased by NAM.** Organotypic retinal explants derived from wild-type (WT) and *Rho*$^{I255d/+}$ at post-natal (P) day 12, treated with HDAC inhibitors (SAHA; S, MPT; M, ACY; A, NAM; N), ended at P20. **A-H**. HDAC activity (HDAC*, white;) was detected *in situ* in the outer nuclear layer (ONL) of both WT and *Rho*$^{I255d/+}$ retina. Quantification of HDAC* positive cells in the ONL revealed increased numbers in non-treated (NT) *Rho*$^{I255d/+}$ retina, compared with WT. **A-B.** SAHA treatment reduced the numbers of HDAC*-positive cells in the ONL at 0.01 and 0.1 µM. **C-D**. MPT decreased HDAC*-positive cells at 0.1 and 1 µM. **E-F**. ACY inhibited HDAC* at 0.01, 0.1, and 1 µM. **G-H**. The sirtuin inhibitor NAM increased overall HDAC* at 1000 µM. ToPro3 was used as nuclear counterstain. Images represent results obtained from 3-6 independent retinal tissues. The strong white label at the image top relates to the culturing membrane; data represent mean and SD; statistical analysis: Two-way ANOVA test with two-stage linear step-up procedure of Benjamini, Krieger, and Yekutieli; * = $p \leqslant 0.05$; ** = $p \leqslant 0.01$; *** = $p \leqslant 0.001$, **** = $p \leqslant 0.0001$.

**Figure S2**

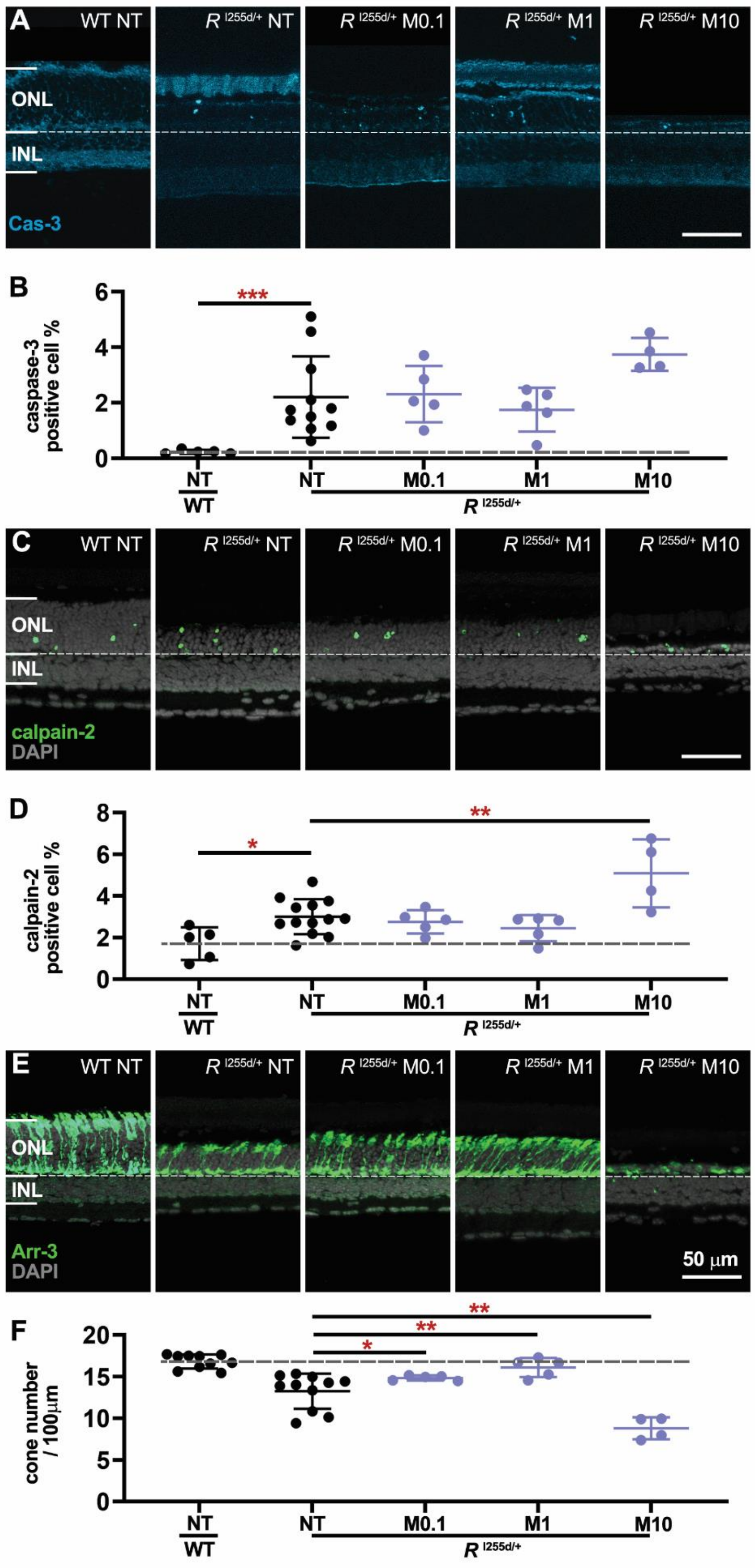

A
WT NT
R I255d/+ NT
R I255d/+ M0.1
R I255d/+ M1
R I255d/+ M10
ONL
INL
Cas-3
B
caspase-3 positive cell %
***
NT
WT
NT
M0.1
M1
M10
R I255d/+
C
calpain-2
DAPI
D
calpain-2 positive cell %
*
**
E
Arr-3
DAPI
50 μm
F
cone number / 100μm

**Figure S2. Differential effects of MPT treatment on caspase-3 and calpain-2 activation, neuroprotection of cones.** Organotypic retinal explants derived from wild-type (WT) and *Rho*$^{I255d/+}$ mice at post-natal (P) day 12 were treated with MPT (M) from P14 to P20**.** **A-B.** Apoptosis was visualized by immunostaining of cleaved caspase-3 (blue, cas-3) in the outer nuclear layer (ONL). Compared to non-treated (NT) WT, the numbers of cas-3 positive cells were increased in NT *Rho*$^{I255d/+}$ mutant. **C-D**. Calpain-2 activation (green) was increased in NT mutant retinas. At 10 µM, MPT treatment increased the number of calpain-2 activation-positive cells. **E-F**. Arrestin-3 (green, Arr-3) labeled cone photoreceptors. Treatment with 0.1 and 1 µM MPT led to significant cone preservation, while at 10 µM cone numbers were decreased. DAPI (grey) was used as the nuclear counterstain. Data represent mean and SD, and were obtained from 4-14 independent retinal explant cultures; statistical analysis: Two-way ANOVA test with two-stage linear step-up procedure of Benjamini, Krieger, and Yekutieli; * = $p \leqslant 0.05$, ** = $p \leqslant 0.01$, *** = $p \leqslant 0.001$.

**Figure S3**

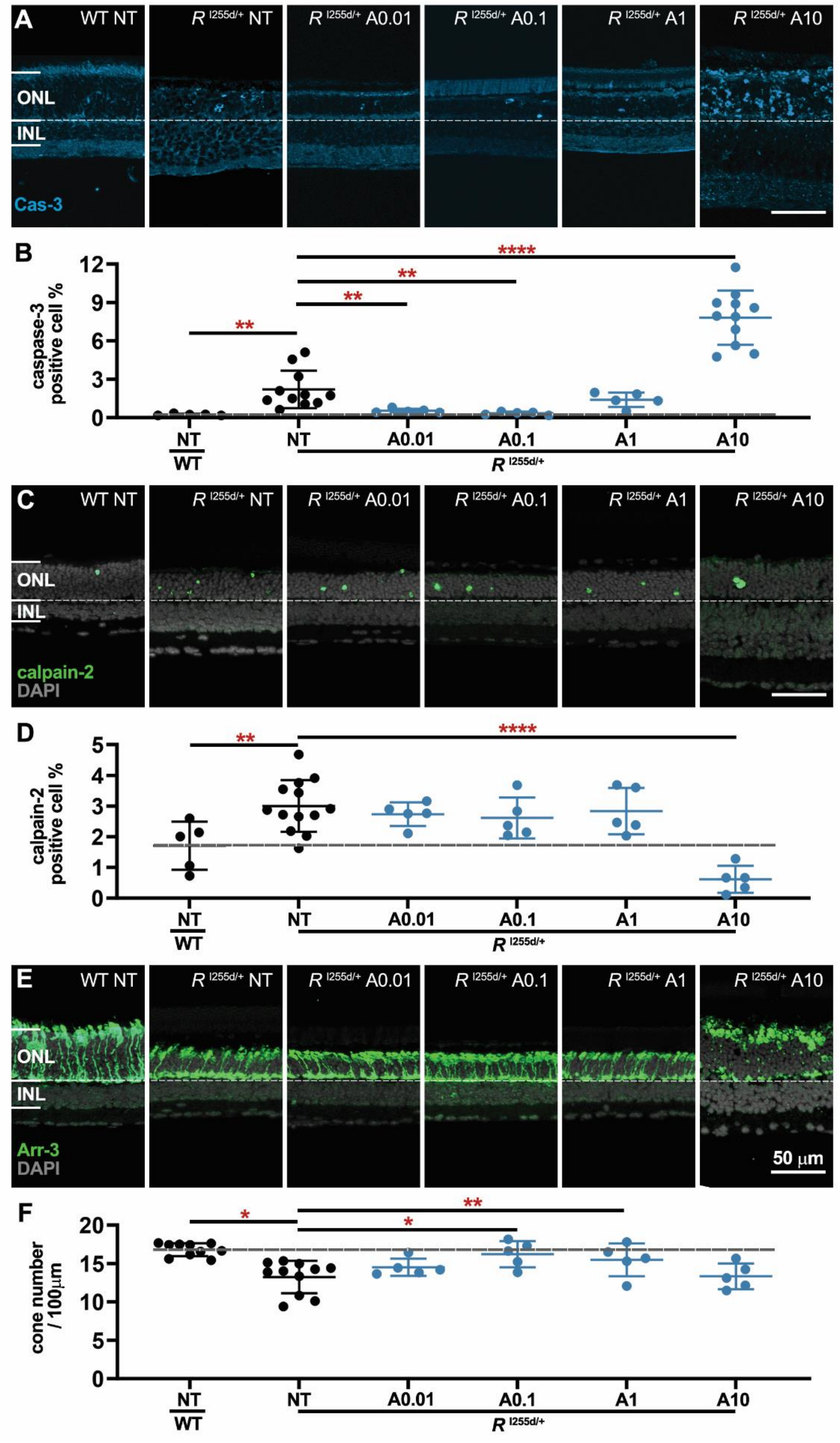

A
WT NT
R I255d/+ NT
R I255d/+ A0.01
R I255d/+ A0.1
R I255d/+ A1
R I255d/+ A10
ONL
INL
Cas-3
B
caspase-3 positive cell %
12
9
6
3
0
**
**
**
****
NT
WT
NT
A0.01
A0.1
A1
A10
R I255d/+
C
calpain-2
DAPI
D
calpain-2 positive cell %
5
4
3
2
1
0
**
****
E
Arr-3
DAPI
50 μm
F
cone number / 100μm
20
15
10
5
0
*
**
*

**Figure S3. ACY differentially affects caspase-3 and calpain-2 activation, preserves cones at low concentrations.** Retinal explants were derived from wild-type (WT) and *Rho*$^{I255d/+}$ mice cultured from post-natal (P) day 12 to P20 and treated with ACY (A) from P14. **A-B.** Apoptosis was displayed by cleaved caspase-3 (blue, cas-3) in the outer nuclear layer (ONL). Non-treated (NT) mutant increased strongly compared with NT WT. The percentage of caspase-3 positive cells in the mutant ONL decreased at lower concentrations of ACY (0.01 and 0.1 µM), but increased at 10 µM. **C-D**. The percentage of calpain-2 (green) positive cells increased in NT mutant compared to NT WT, but decreased at 10 µM in the ONL of *Rho*$^{I255d/+}$ compared to NT. **E-F.** Arrestin-3 (green, Arr-3) labelled cone photoreceptors and their number was reduced in the NT mutant, compared to WT. Remarkably, treatment of ACY at 0.1 and 1 µM showed neuroprotection of cone photoreceptors. However, the ONL structure appeared disrupted at 10 µM. Nuclear counterstain by DAPI (dark grey). Data represent mean and SD, and were obtained from 5-13 independent retinal explant cultures; statistical analysis: Two-way ANOVA test with two-stage linear step-up procedure of Benjamini, Krieger, and Yekutieli; * = $p \leqslant 0.05$, ** = $p \leqslant 0.01$, **** = $p \leqslant 0.0001$.

**Figure S4**

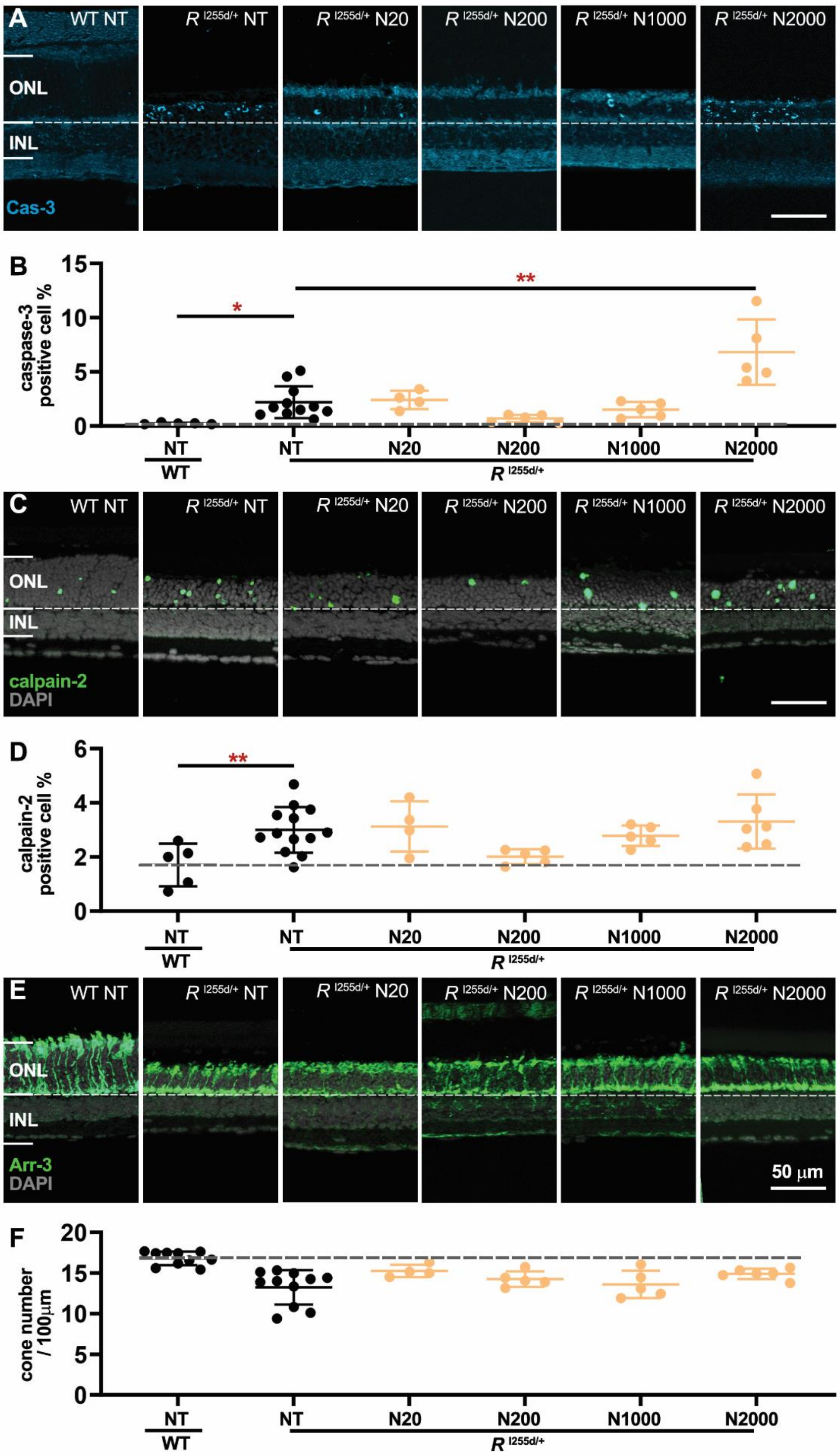

A
WT NT
$R^{I255d/+}$ NT
$R^{I255d/+}$ N20
$R^{I255d/+}$ N200
$R^{I255d/+}$ N1000
$R^{I255d/+}$ N2000
ONL
INL
Cas-3
B
caspase-3 positive cell %
15
10
5
0
*
**
NT
WT
NT
N20
N200
N1000
N2000
$R^{I255d/+}$
C
calpain-2
DAPI
D
calpain-2 positive cell %
6
4
2
0
**
E
Arr-3
50 μm
F
cone number / 100μm
20
15
10
5
0

**Figure S4. NAM causes *Rho*$^{I255d/+}$ rod degeneration, without affecting cones.** Organotypic retinal explants were obtained from wild-type (WT) and *Rho*$^{I255d/+}$ mice at postnatal day 12 (P12), and were treated with NAM from P14 to P20. **A-B.** Cleaved caspase-3 (blue, cas-3) in the outer nuclear layer (ONL) represented apoptosis. The value of caspase-3 positive cells increased in non-treated (NT) *Rho*$^{I255d/+}$ compared with WT, and it was stimulated under treatment with NAM at 2000 µM. **C-D**. Calpain-2 (green) activation increased in NT *Rho*$^{I255d/+}$ compared to WT, but 200 µM NAM decreased calpain-2 activation. **E-F.** The number of arrestin-3 (green, Arr-3) positive cone photoreceptors remained unaffected under treatment. DAPI (dark grey) was used as a nuclear counterstain. Images represent results from 5-13 independent retinal explant cultures; data represent mean and SD; statistical analysis: Two-way ANOVA test with two-stage linear step-up procedure of Benjamini, Krieger, and Yekutieli; * = $p \leqslant 0.05$, ** = $p \leqslant 0.01$.

### Figure S5

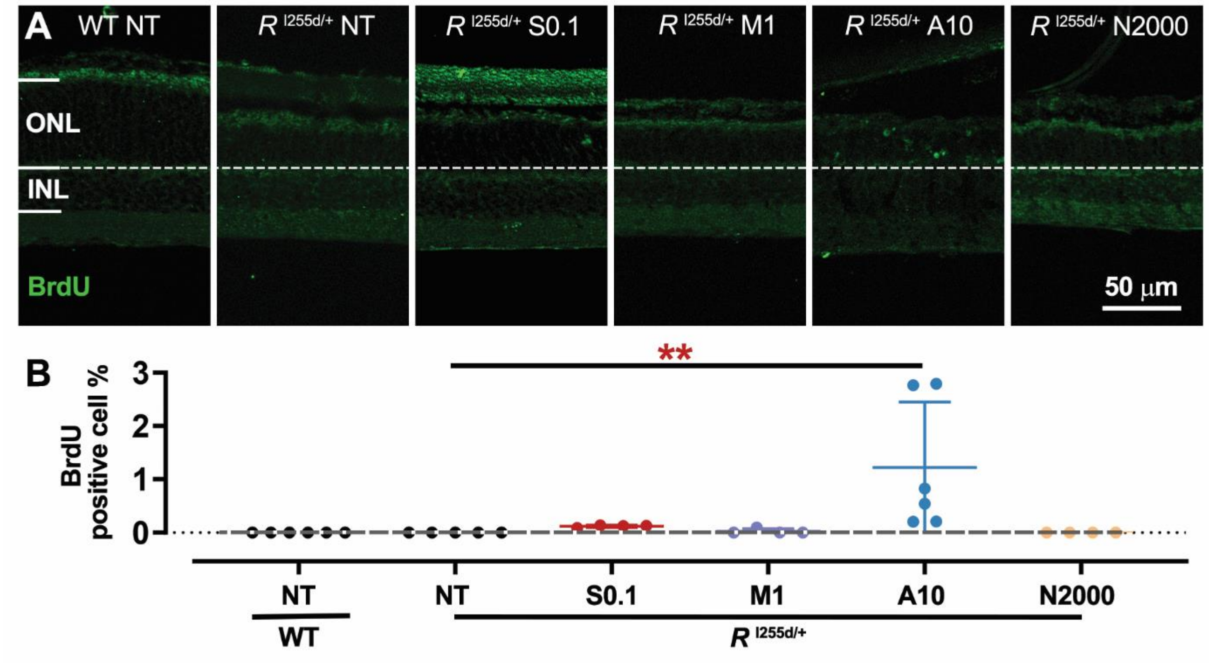


**Figure S5. ACY stimulates cell proliferation in *Rho*$^{I255d/+}$ outer nuclear layer.** Retinal explants derived from wild-type (WT) or *Rho*$^{I255d/+}$ mutant were treated with HDAC inhibitors SAHA (S; 0.1 µM), MPT (M; 1 µM), ACY (A; 10 µM), or NAM (N; 2000 µM) from post-natal (P) day 14 to 20. From P16 to P18, these cultures were exposed to the cellular proliferation marker bromodeoxyuridine (BrdU) and then stained with an antibody directed against BrdU (green) at P20. **A**. Positive BrdU labeling was seen in the outer nuclear layer (ONL) of retinas treated with ACY. **B.** Data plot showing the percentage of BrdU-positive cells in the ONL. ACY treatment increased BrdU in the mutant retina. Images represent results obtained from 5-6 independent retinal explant cultures; data points represent mean and SD; statistical analysis: Two-way ANOVA test with two-stage linear step-up procedure of Benjamini, Krieger, and Yekutieli; ** = $p \leqslant 0.01$.
