## supplemental tables for "Differential effects of HDAC inhibitors in the *Rho*^I255d^ mouse model for autosomal dominant retinitis pigmentosa"

**Table S1.** Increased HDAC activity in *Rho*$^{I255d/+}$ photoreceptors and application of HDAC inhibitors (TSA and NAM): Quantitative data for graphs presented in Figure 1E.

| Table S1 (Figure) | Genotype | *p* - value comparison | Genotype-treatment | Mean ± SD (%) | *p* - value | n |
|---|---|---|---|---|---|---|
| Fig. 1E | WT | *R*$^{I255d/+}$ - NT | NT | 0.44 ± 0.34 | *p* < 0.0001 | 6 |
| | | | TSA | 0.24 ± 0.30 | *p* < 0.0001 | 3 |
| | | | NAM | 0.32 ± 0.35 | *p* < 0.0001 | 3 |
| | | | TSA+NAM | 0.19 ± 0.21 | *p* < 0.0001 | 3 |
| | *R*$^{I255d/+}$ | *R*$^{I255d/+}$ - NT | NT | 3.55 ± 1.39 | | 6 |
| | | | TSA | 1.26 ± 0.30 | *p* < 0.0001 | 6 |
| | | | NAM | 1.97 ± 0.89 | *p* = 0.0003 | 6 |
| | | | TSA+NAM | 0.94 ± 0.52 | *p* < 0.0001 | 6 |
| | | *R*$^{I255d/+}$ - TSA | NAM | | *p* = 0.0426 | |
| | | *R*$^{I255d/+}$ - TSA | TSA+NAM | | *p* = 0.3475 | |
| | | *R*$^{I255d/+}$ - NAM | TSA+NAM | | *p* = 0.0049 | |

**Table S2.** Effect of SAHA (S) on cell death (TUNEL assay) in outer nuclear layer (ONL) and inner nuclear layer (INL), and ONL row counts, with concentrations (0.01, 0.1, and 1 µM): Quantitative data for graphs presented in Figures 2B, C, D.

| Table S2 (Figure) | Parameter | *p* - value comparison | Genotype-treatment | Mean ± SD (%) | *p* - value | n |
|---|---|---|---|---|---|---|
| Fig. 2B | ONL-TUNEL | WT - NT | WT - NT | 2.06 ± 0.87 | | 5 |
| | | | WT - S 0.01 | 2.91 ± 0.38 | *p* = 0.3227 | 6 |
| | | | WT - S 0.1 | 2.76 ± 0.48 | *p* = 0.3409 | 4 |
| | | | WT - S 1 | 9.57 ± 2.54 | *p* < 0.0001 | 3 |
| | | $R^{I255d/+}$ - NT | WT - NT | | *p* < 0.0001 | |
| | | | $R^{I255d/+}$ - NT | 7.29 ± 1.87 | | 14 |
| | | | $R^{I255d/+}$ - S 0.01 | 7.94 ± 0.90 | *p* = 0.3989 | 5 |
| | | | $R^{I255d/+}$ - S 0.1 | 3.44 ± 1.50 | *p* < 0.0001 | 10 |
| | | | $R^{I255d/+}$ - S 1 | 7.18 ± 1.48 | *p* = 0.4344 | 5 |
| Fig. 2C | INL-TUNEL | WT - NT | WT - NT | 0.09 ± 0.11 | | 5 |
| | | | WT - S 0.01 | 0.09 ± 0.09 | *p* = 0.9929 | 6 |
| | | | WT - S 0.1 | 0.08 ± 0.09 | *p* = 0.8274 | 6 |
| | | | WT - S 1 | 0.05 ± 0.09 | *p* = 0.8646 | 3 |
| | | $R^{I255d/+}$ - NT | WT - NT | | *p* = 0.3265 | |
| | | | $R^{I255d/+}$ - NT | 0.09 ± 0.10 | | 14 |
| | | | $R^{I255d/+}$ - S 0.01 | 0.07 ± 0.03 | *p* = 0.1660 | 5 |
| | | | $R^{I255d/+}$ - S 0.1 | 0.10 ± 0.09 | *p* = 0.9495 | 10 |
| | | | $R^{I255d/+}$ - S 1 | 0.14 ± 0.16 | *p* = 0.9880 | 5 |
| Fig. 2D | ONL rows | WT - NT | WT - NT | 7.52 ± 0.43 | | 5 |
| | | | WT - S 0.01 | 7.87 ± 0.30 | *p* = 0.4872 | 6 |
| | | | WT - S 0.1 | 7.98 ± 0.84 | *p* = 0.3863 | 4 |
| | | | WT - S 1 | 5.98 ± 0.74 | *p* = 0.0104 | 3 |
| | | $R^{I255d/+}$ - NT | WT - NT | | *p* < 0.0001 | |
| | | | $R^{I255d/+}$ - NT | 4.99 ± 0.60 | | 14 |
| | | | $R^{I255d/+}$ - S 0.01 | 4.19 ± 0.47 | *p* = 0.0345 | 5 |
| | | | $R^{I255d/+}$ - S 0.1 | 5.14 ± 0.74 | *p* = 0.2990 | 6 |
| | | | $R^{I255d/+}$ - S 1 | 1.51 ± 0.30 | *p* < 0.0001 | 5 |

**Table S3.** Effect of SAHA (S) on caspase-3 (cleaved-caspase-3 staining), calpain-2 (calpain-2 staining) and cone survival (cone arrestin-3 staining) in outer nuclear layer (ONL), with concentrations (0.01, 0.1, and 1 µM): Quantitative data for graphs presented in Figures 3B, D, F.

| Table S3 (Figure) | Parameter | *p* - value comparison | Genotype-treatment | Mean ± SD (%) | *p* - value | n |
|---|---|---|---|---|---|---|
| Fig. 3B | Caspase-3 | WT - NT | WT - NT | 0.23 ± 0.07 | | 5 |
| | | | WT - S 0.01 | 0.22 ± 0.07 | $p = 0.3227$ | 6 |
| | | | WT - S 0.1 | 0.42 ± 0.24 | $p = 0.3409$ | 3 |
| | | | WT - S 1 | 0.38 ± 0.36 | $p < 0.0001$ | 3 |
| | | $R^{I255d/+}$ - NT | WT - NT | | $p = 0.0002$ | |
| | | | $R^{I255d/+}$ - NT | 2.20 ± 1.47 | | 11 |
| | | | $R^{I255d/+}$ - S 0.01 | 2.04 ± 0.49 | $p = 0.2133$ | 5 |
| | | | $R^{I255d/+}$ - S 0.1 | 0.47 ± 0.36 | $p = 0.0003$ | 9 |
| | | | $R^{I255d/+}$ - S 1 | 0.12 ± 0.09 | $p = 0.0004$ | 4 |
| Fig. 3D | Calpain-2 | WT - NT | WT - NT | 1.71 ± 0.78 | | 5 |
| | | | WT - S 0.01 | 2.12 ± 0.38 | $p = 0.3558$ | 6 |
| | | | WT - S 0.1 | 2.24 ± 0.11 | $p = 0.0756$ | 3 |
| | | | WT - S 1 | 3.84 ± 0.71 | $p = 0.0004$ | 3 |
| | | $R^{I255d/+}$ - NT | WT - NT | | $p = 0.0347$ | |
| | | | $R^{I255d/+}$ - NT | 3.00 ± 0.84 | | 13 |
| | | | $R^{I255d/+}$ - S 0.01 | 2.94 ± 0.30 | $p = 0.8061$ | 5 |
| | | | $R^{I255d/+}$ - S 0.1 | 1.64 ± 0.58 | $p = 0.0020$ | 9 |
| | | | $R^{I255d/+}$ - S 1 | 2.10 ± 1.04 | $p = 0.1574$ | 5 |
| Fig. 3F | Cone | WT - NT | WT - NT | 16.81 ± 0.85 | | 10 |
| | | | WT - S 0.01 | 15.56 ± 1.29 | $p = 0.0186$ | 6 |
| | | | WT - S 0.1 | 15.34 ± 0.82 | $p = 0.9559$ | 3 |
| | | | WT - S 1 | 11.83 ± 0.30 | $p = 0.0032$ | 3 |
| | | $R^{I255d/+}$ - NT | WT - NT | | $p = 0.0600$ | |
| | | | $R^{I255d/+}$ - NT | 13.24 ± 2.11 | | 11 |
| | | | $R^{I255d/+}$ - S 0.01 | 14.72 ± 0.85 | $p = 0.6715$ | 5 |
| | | | $R^{I255d/+}$ - S 0.1 | 12.99 ± 2.61 | $p = 0.6423$ | 7 |
| | | | $R^{I255d/+}$ - S 1 | 7.56 ± 0.99 | $p = 0.0002$ | 4 |

**Table S4.** Effect of MPT (M) on cell death (TUNEL assay) in outer nuclear layer (ONL) and inner nuclear layer (INL) and ONL row counts, with concentrations (0.1, 1, and 10 µM): Quantitative data for graphs presented in Figures 4B, C, D.

| Table S4 (Figure) | Parameter | *p* - value comparison | Genotype-treatment | Mean ± SD (%) | *p* - value | n |
|---|---|---|---|---|---|---|
| Fig. 4B | ONL-TUNEL | $R^{I255d/+}$ - NT | WT - NT | 2.06 ± 0.87 | $p = 0.0002$ | 5 |
| | | | $R^{I255d/+}$ - NT | 7.29 ± 1.87 | | 14 |
| | | | $R^{I255d/+}$ - M 0.1 | 4.94 ± 1.44 | $p = 0.0711$ | 5 |
| | | | $R^{I255d/+}$ - M 1 | 3.60 ± 0.67 | $p = 0.0045$ | 5 |
| | | | $R^{I255d/+}$ - M 10 | 9.80 ± 1.69 | $p = 0.0113$ | 4 |
| Fig. 4C | INL-TUNEL | $R^{I255d/+}$ - NT | WT - NT | 0.09 ± 0.11 | $p = 0.2856$ | 5 |
| | | | $R^{I255d/+}$ - NT | 0.09 ± 0.10 | | 14 |
| | | | $R^{I255d/+}$ - M 0.1 | 0.07 ± 0.08 | $p = 0.1676$ | 5 |
| | | | $R^{I255d/+}$ - M 1 | 0.04 ± 0.02 | $p = 0.0468$ | 5 |
| | | | $R^{I255d/+}$ - M 10 | 0.14 ± 0.08 | $p = 0.7974$ | 4 |
| Fig. 4D | ONL rows | $R^{I255d/+}$ - NT | WT - NT | 7.52 ± 0.43 | $p < 0.0001$ | 5 |
| | | | $R^{I255d/+}$ - NT | 4.99 ± 0.60 | | 14 |
| | | | $R^{I255d/+}$ - M 0.1 | 5.49 ± 0.86 | $p = 0.0692$ | 5 |
| | | | $R^{I255d/+}$ - M 1 | 5.41 ± 0.62 | $p = 0.1084$ | 5 |
| | | | $R^{I255d/+}$ - M 10 | 1.23 ± 0.31 | $p < 0.0001$ | 4 |

**Table S5.** Effect of ACY (A) on cell death (TUNEL assay) in outer nuclear layer (ONL) and inner nuclear layer (INL) and ONL row counts, with concentrations (0.01, 0.1, 1, and 10 µM): Quantitative data for graphs presented in Figures 5B, C, D.

| Table S5 (Figure) | Parameter | *p* - value comparison | Genotype-treatment | Mean ± SD (%) | *p* - value | n |
|---|---|---|---|---|---|---|
| Fig. 5B | ONL-TUNEL | $R^{I255d/+}$ - NT | WT - NT | 2.06 ± 0.87 | $p = 0.0472$ | 5 |
| | | | $R^{I255d/+}$ - NT | 7.29 ± 1.87 | | 14 |
| | | | $R^{I255d/+}$ - A 0.01 | 5.63 ± 1.00 | $p = 0.6049$ | 5 |
| | | | $R^{I255d/+}$ - A 0.1 | 5.20 ± 0.66 | $p = 0.4823$ | 5 |
| | | | $R^{I255d/+}$ - A 1 | 4.45 ± 1.06 | $p = 0.3050$ | 5 |
| | | | $R^{I255d/+}$ - A 10 | 25.09 ± 7.91 | $p < 0.0001$ | 5 |
| Fig. 5C | INL-TUNEL | $R^{I255d/+}$ - NT | WT - NT | 0.09 ± 0.11 | $p = 0.7302$ | 5 |
| | | | $R^{I255d/+}$ - NT | 0.09 ± 0.10 | | 14 |
| | | | $R^{I255d/+}$ - A 0.01 | 0.07 ± 0.05 | $p = 0.6187$ | 5 |
| | | | $R^{I255d/+}$ - A 0.1 | 0.01 ± 0.02 | $p = 0.3894$ | 5 |
| | | | $R^{I255d/+}$ - A 1 | 0.00 ± 0.00 | $p = 0.3538$ | 5 |
| | | | $R^{I255d/+}$ - A 10 | 1.46 ± 0.55 | $p < 0.0001$ | 5 |
| Fig. 5D | ONL rows | $R^{I255d/+}$ - NT | WT - NT | 7.52 ± 0.43 | $p < 0.0001$ | 5 |
| | | | $R^{I255d/+}$ - NT | 4.99 ± 0.60 | | 14 |
| | | | $R^{I255d/+}$ - A 0.01 | 5.20 ± 0.76 | $p = 0.3688$ | 5 |
| | | | $R^{I255d/+}$ - A 0.1 | 5.21 ± 0.25 | $p = 0.3504$ | 5 |
| | | | $R^{I255d/+}$ - A 1 | 5.15 ± 0.47 | $p = 0.4489$ | 5 |
| | | | $R^{I255d/+}$ - A 10 | 5.84 ± 0.82 | $p = 0.0142$ | 5 |

**Table S6.** Effect of NAM (N) on cell death (TUNEL assay) in outer nuclear layer (ONL) and inner nuclear layer (INL) and ONL row counts, with concentrations (20, 200, 1000, and 2000 µM): Quantitative data for graphs presented in Figures 6B, C, D.

| Table S6 (Figure) | Parameter | *p* - value comparison | Genotype-treatment | Mean ± SD (%) | *p* - value | n |
|---|---|---|---|---|---|---|
| Fig. 6B | ONL-TUNEL | $R^{I255d/+}$ - NT | WT - NT | 2.06 ± 0.87 | $p = 0.0002$ | 5 |
| | | | $R^{I255d/+}$ - NT | 7.29 ± 1.87 | | 14 |
| | | | $R^{I255d/+}$ - N 20 | 9.74 ± 1.89 | $p = 0.2577$ | 4 |
| | | | $R^{I255d/+}$ - N 200 | 11.97 ± 2.66 | $p = 0.0031$ | 5 |
| | | | $R^{I255d/+}$ - N 1000 | 15.79 ± 1.29 | $p < 0.0001$ | 5 |
| | | | $R^{I255d/+}$ - N 2000 | 16.37 ± 1.74 | $p < 0.0001$ | 7 |
| Fig. 6C | INL-TUNEL | $R^{I255d/+}$ - NT | WT - NT | 0.09 ± 0.11 | $p = 0.4600$ | 5 |
| | | | $R^{I255d/+}$ - NT | 0.09 ± 0.10 | | 14 |
| | | | $R^{I255d/+}$ - N 20 | 0.02 ± 0.05 | $p = 0.1123$ | 4 |
| | | | $R^{I255d/+}$ - N 200 | 0.06 ± 0.06 | $p = 0.1944$ | 5 |
| | | | $R^{I255d/+}$ - N 1000 | 0.12 ± 0.11 | $p = 0.8939$ | 5 |
| | | | $R^{I255d/+}$ - N 2000 | 0.01 ± 0.02 | $p = 0.0274$ | 5 |
| Fig. 6D | ONL rows | $R^{I255d/+}$ - NT | WT - NT | 7.52 ± 0.43 | $p < 0.0001$ | 5 |
| | | | $R^{I255d/+}$ - NT | 4.99 ± 0.60 | | 14 |
| | | | $R^{I255d/+}$ - N 20 | 5.32 ± 0.94 | $p = 0.2740$ | 4 |
| | | | $R^{I255d/+}$ - N 200 | 5.39 ± 0.26 | $p = 0.1929$ | 5 |
| | | | $R^{I255d/+}$ - N 1000 | 5.41 ± 0.63 | $p = 0.1721$ | 5 |
| | | | $R^{I255d/+}$ - N 2000 | 5.37 ± 0.34 | $p = 0.2041$ | 5 |

**Table S7.** Effect of different HDAC inhibitors on HDAC activity (HDAC activity *in situ* assay) in outer nuclear layer (ONL). Treatments with SAHA (S) at 0.01 and 0.1 µM, MPT (M) at 0.1 and 1 µM, ACY (A) at 0.01, 0.1 and 1 µM, and NAM (N) at 20, 200, 1000, and 2000 µM: Quantitative data for graphs presented in Figures S1B, D, F, H.

| Table S7 (Figure) | Treatment | *p* - value comparison | Genotype-treatment | Mean ± SD (%) | *p* - value | n |
|---|---|---|---|---|---|---|
| Fig. S1B | SAHA | $R^{I255d/+}$ - NT | WT - NT | 0.42 ± 0.09 | $p = 0.0009$ | 3 |
| | | | $R^{I255d/+}$ - NT | 2.33 ± 0.78 | | 5 |
| | | | $R^{I255d/+}$ - S 0.01 | 1.13 ± 0.32 | $p = 0.0067$ | 5 |
| | | | $R^{I255d/+}$ - S 0.1 | 0.71 ± 0.23 | $p = 0.0015$ | 6 |
| Fig. S1D | MPT | $R^{I255d/+}$ - NT | WT - NT | 0.42 ± 0.09 | $p < 0.0001$ | 3 |
| | | | $R^{I255d/+}$ - NT | 2.33 ± 0.78 | | 5 |
| | | | $R^{I255d/+}$ - M 0.1 | 1.61 ± 0.27 | $p = 0.0237$ | 5 |
| | | | $R^{I255d/+}$ - M 1 | 1.72 ± 0.40 | $p = 0.0483$ | 5 |
| Fig. S1F | ACY | $R^{I255d/+}$ - NT | WT - NT | 0.42 ± 0.09 | $p < 0.0001$ | 3 |
| | | | $R^{I255d/+}$ - NT | 2.33 ± 0.78 | | 5 |
| | | | $R^{I255d/+}$ - A 0.01 | 1.48 ± 0.24 | $p = 0.0049$ | 5 |
| | | | $R^{I255d/+}$ - A 0.1 | 1.03 ± 0.26 | $p = 0.0002$ | 5 |
| | | | $R^{I255d/+}$ - A 1 | 0.79 ± 0.19 | $p < 0.0001$ | 5 |
| Fig. S1H | NAM | $R^{I255d/+}$ - NT | WT - NT | 0.42 ± 0.09 | $p = 0.0396$ | 3 |
| | | | $R^{I255d/+}$ - NT | 2.33 ± 0.78 | | 5 |
| | | | $R^{I255d/+}$ - N 20 | 1.39 ± 0.40 | $p = 0.3438$ | 4 |
| | | | $R^{I255d/+}$ - N 200 | 3.29 ± 0.75 | $p = 0.1347$ | 5 |
| | | | $R^{I255d/+}$ - N 1000 | 4.04 ± 0.84 | $p = 0.0125$ | 5 |
| | | | $R^{I255d/+}$ - N 2000 | 4.36 ± 3.09 | $p = 0.1168$ | 6 |

**Table S8.** Effect of MPT (M) on caspase-3 (cleaved-caspase-3 staining), calpain-2 (calpain-2 staining) and cone survival (cone arrestin-3 staining) in outer nuclear layer (ONL), with concentrations (0.1, 1, and 10 μM): Quantitative data for graphs presented in Figures S2B, D, F.

| Table S8 (Figure) | Parameter | *p* - value comparison | Genotype-treatment | Mean ± SD (%) | *p* - value | n |
|---|---|---|---|---|---|---|
| Fig. S2B | Caspase-3 | $R^{I255d/+}$ - NT | WT - NT | 0.23 ± 0.07 | *p* = 0.0008 | 5 |
| | | | $R^{I255d/+}$ - NT | 2.20 ± 1.47 | | 11 |
| | | | $R^{I255d/+}$ - M 0.1 | 2.31 ± 1.02 | *p* = 0.2620 | 5 |
| | | | $R^{I255d/+}$ - M 1 | 1.75 ± 0.79 | *p* = 0.0661 | 5 |
| | | | $R^{I255d/+}$ - M 10 | 3.74 ± 0.59 | *p* = 0.4677 | 4 |
| Fig. S2D | Calpain-2 | $R^{I255d/+}$ - NT | WT - NT | 1.71 ± 0.78 | *p* = 0.0361 | 5 |
| | | | $R^{I255d/+}$ - NT | 3.00 ± 0.84 | | 13 |
| | | | $R^{I255d/+}$ - M 0.1 | 2.75 ± 0.56 | *p* = 0.6803 | 5 |
| | | | $R^{I255d/+}$ - M 1 | 2.45 ± 0.62 | *p* = 0.3514 | 5 |
| | | | $R^{I255d/+}$ - M 10 | 5.08 ± 1.63 | *p* = 0.0035 | 4 |
| Fig. S2F | Cone | $R^{I255d/+}$ - NT | WT - NT | 16.81 ± 0.85 | *p* = 0.1148 | 10 |
| | | | $R^{I255d/+}$ - NT | 13.24 ± 2.11 | | 11 |
| | | | $R^{I255d/+}$ - M 0.1 | 14.83 ± 0.31 | *p* = 0.0186 | 5 |
| | | | $R^{I255d/+}$ - M 1 | 16.09 ± 1.14 | *p* = 0.0018 | 5 |
| | | | $R^{I255d/+}$ - M 10 | 8.79 ± 1.33 | *p* = 0.0051 | 4 |

**Table S9.** Effect of ACY (A) on caspase-3 (cleaved-caspase-3 staining), calpain-2 (calpain-2 staining) and cone survival (cone arrestin-3 staining) in outer nuclear layer (ONL), with concentrations (0.01, 0.1, 1, and 10 µM): Quantitative data for graphs presented in Figures S3B, D, F.

| Table S9 (Figure) | Parameter | *p* - value comparison | Genotype-treatment | Mean ± SD (%) | *p* - value | n |
|---|---|---|---|---|---|---|
| Fig. S3B | Caspase-3 | $R^{\mathrm{I255d/+}}$ - NT | WT - NT | 0.23 ± 0.07 | $p = 0.0018$ | 5 |
| | | | $R^{\mathrm{I255d/+}}$ - NT | 2.20 ± 1.47 | | 11 |
| | | | $R^{\mathrm{I255d/+}}$ - A 0.01 | 0.54 ± 0.16 | $p = 0.0046$ | 5 |
| | | | $R^{\mathrm{I255d/+}}$ - A 0.1 | 0.34 ± 0.12 | $p = 0.0025$ | 5 |
| | | | $R^{\mathrm{I255d/+}}$ - A 1 | 1.40 ± 0.57 | $p = 0.0537$ | 5 |
| | | | $R^{\mathrm{I255d/+}}$ - A 10 | 7.81 ± 2.12 | $p < 0.0001$ | 11 |
| Fig. S3D | Calpain-2 | $R^{\mathrm{I255d/+}}$ - NT | WT - NT | 1.71 ± 0.78 | $p = 0.0027$ | 5 |
| | | | $R^{\mathrm{I255d/+}}$ - NT | 3.00 ± 0.84 | | 13 |
| | | | $R^{\mathrm{I255d/+}}$ - A 0.01 | 2.74 ± 0.39 | $p = 0.5056$ | 5 |
| | | | $R^{\mathrm{I255d/+}}$ - A 0.1 | 2.61 ± 0.67 | $p = 0.3285$ | 5 |
| | | | $R^{\mathrm{I255d/+}}$ - A 1 | 2.83 ± 0.76 | $p = 0.6846$ | 5 |
| | | | $R^{\mathrm{I255d/+}}$ - A 10 | 0.61 ± 0.44 | $p < 0.0001$ | 5 |
| Fig. S3F | Cone | $R^{\mathrm{I255d/+}}$ - NT | WT - NT | 16.81 ± 0.85 | $p = 0.0383$ | 10 |
| | | | $R^{\mathrm{I255d/+}}$ - NT | 13.24 ± 2.11 | | 11 |
| | | | $R^{\mathrm{I255d/+}}$ - A 0.01 | 14.51 ± 1.12 | $p = 0.0508$ | 5 |
| | | | $R^{\mathrm{I255d/+}}$ - A 0.1 | 16.22 ± 1.71 | $p = 0.0414$ | 5 |
| | | | $R^{\mathrm{I255d/+}}$ - A 1 | 15.48 ± 2.14 | $p = 0.0083$ | 5 |
| | | | $R^{\mathrm{I255d/+}}$ - A 10 | 13.34 ± 1.67 | $p = 0.3228$ | 5 |

**Table S10.** Effect of NAM (N) on caspase-3 (cleaved-caspase-3 staining), calpain-2 (calpain-2 staining) and cone survival (cone arrestin-3 staining) in outer nuclear layer (ONL), with concentrations (20, 200, 1000, and 2000 µM): Quantitative data for graphs presented in Figures S4B, D, F.

| Table S10 (Figure) | Parameter | *p* - value comparison | Genotype-treatment | Mean ± SD (%) | *p* - value | n |
|---|---|---|---|---|---|---|
| Fig. S4B | Caspase-3 | $R^{I255d/+}$ - NT | WT - NT | 0.23 ± 0.07 | *p* = 0.0257 | 5 |
| | | | $R^{I255d/+}$ - NT | 2.20 ± 1.47 | | 11 |
| | | | $R^{I255d/+}$ - N 20 | 2.41 ± 0.84 | *p* = 0.8448 | 4 |
| | | | $R^{I255d/+}$ - N 200 | 0.70 ± 0.34 | *p* = 0.1702 | 5 |
| | | | $R^{I255d/+}$ - N 1000 | 1.52 ± 0.70 | *p* = 0.4835 | 5 |
| | | | $R^{I255d/+}$ - N 2000 | 6.83 ± 3.02 | *p* = 0.0027 | 5 |
| Fig. S4D | Calpain-2 | $R^{I255d/+}$ - NT | WT - NT | 1.71 ± 0.78 | *p* = 0.0069 | 5 |
| | | | $R^{I255d/+}$ - NT | 3.00 ± 0.84 | | 13 |
| | | | $R^{I255d/+}$ - N 20 | 3.13 ± 0.93 | *p* = 0.9456 | 4 |
| | | | $R^{I255d/+}$ - N 200 | 1.94 ± 0.05 | *p* = 0.3294 | 5 |
| | | | $R^{I255d/+}$ - N 1000 | 2.79 ± 0.38 | *p* = 0.5698 | 5 |
| | | | $R^{I255d/+}$ - N 2000 | 3.31 ± 1.00 | *p* = 0.1491 | 6 |
| Fig. S4F | Cone | $R^{I255d/+}$ - NT | WT - NT | 16.81 ± 0.85 | *p* = 0.0749 | 10 |
| | | | $R^{I255d/+}$ - NT | 13.24 ± 2.11 | | 11 |
| | | | $R^{I255d/+}$ - N 20 | 15.25 ± 0.76 | *p* = 0.5261 | 4 |
| | | | $R^{I255d/+}$ - N 200 | 14.26 ± 0.95 | *p* = 0.6192 | 5 |
| | | | $R^{I255d/+}$ - N 1000 | 13.60 ± 1.67 | *p* = 0.2330 | 5 |
| | | | $R^{I255d/+}$ - N 2000 | 14.91 ± 0.64 | *p* = 0.8426 | 6 |

**Table S11.** Assessment of photoreceptor proliferation using bromodeoxyuridine (BrdU) treatment from post-natal day (P) 16-18, combined with HDAC inhibitors treatments from P14-P20, including SAHA (S) at 0.1 µM, MPT (M) at 1 µM, ACY (A) at 10 µM, and NAM (N) at 2000 µM: Quantitative data for graphs presented in Figure S5B.

| Table S11 (Figure) | Parameter | *p* - value comparison | Genotype-treatment | Mean ± SD (%) | *p* - value | n |
|---|---|---|---|---|---|---|
| Fig. S5B | BrdU | $R^{I255d/+}$ - NT | WT - NT | 0.00 ± 0.00 | *p* = 0.7721 | 6 |
| | | | $R^{I255d/+}$ - NT | 0.00 ± 0.00 | | 5 |
| | | | $R^{I255d/+}$ - S 0.1 | 0.12 ± 0.02 | *p* = 0.9631 | 4 |
| | | | $R^{I255d/+}$ - M 1 | 0.02 ± 0.05 | *p* = 0.8423 | 4 |
| | | | $R^{I255d/+}$ - A 10 | 1.22 ± 1.23 | *p* = 0.0012 | 6 |
| | | | $R^{I255d/+}$ - N 2000 | 0.00 ± 0.00 | *p* = 0.7926 | 4 |
